# A glucose starvation response governs endocytic trafficking and eisosomal retention of surface cargoes

**DOI:** 10.1101/2020.02.05.936187

**Authors:** Kamilla M.E. Laidlaw, Daniel D. Bisinski, Sviatlana Shashkova, Katherine M. Paine, Malaury A. Veillon, Mark C. Leake, Chris MacDonald

## Abstract

Eukaryotic cells adapt their metabolism to the extracellular environment. Downregulation of surface cargo proteins in response to nutrient stress reduces the burden of anabolic processes whilst elevating catabolic production in the lysosome. We use yeast to show that glucose starvation triggers a transcriptional response that simultaneously increases internalisation from the plasma membrane whilst supressing recycling from endosomes back to the surface. Nuclear export of the Mig1 transcriptional repressor in response to glucose starvation increases levels of the Yap1801/02 clathrin adaptors, which is sufficient to increase cargo internalisation. We also show Gpa1, which coordinates endosomal lipid homeostasis, is required for surface recycling of cargo. Nuclear translocation of Mig1 increases *GPA2* levels and inhibits recycling, potentially by diverting Gpa1 to the surface and interfering with its endosomal role in recycling. Finally, we show glucose starvation results in transcriptional upregulation of many eisosomal factors, which serve to sequester a portion of nutrient transporters to persist the starvation period and maximise nutrient uptake upon return to replete conditions. This latter mechanism provides a physiological benefit for cells to rapidly recover from glucose starvation. Collectively, this remodelling of the surface protein landscape during glucose starvation calibrates metabolism to available nutrients.

## INTRODUCTION

Cell surface membrane protein cargoes perform diverse roles, including initiating signal transduction pathways, up-taking nutrients and maintaining the ionic balance of the cell. Surface proteins are controlled by various complex and overlapping trafficking routes, many of which are conserved throughout evolution (Feyder et al., 2015). Endocytosis involves the internalisation of surface proteins from the plasma membrane (PM) in vesicles that are transported to endosomal compartments. Endocytic trafficking from the PM provides a layer of control, where cargoes can either be temporarily removed from the surface and recycled back or permanently removed via their ubiquitin-mediated degradation in the lysosome (MacDonald and Piper, 2016). Endocytic events can be relatively specific, for example the internalisation of a single receptor, or much more general, where a large and varied cohort of proteins are endocytosed. In response to starvation, it is thought that many surface proteins are downregulated as a survival mechanism to reduce energy consumption via non-essential anabolic processes whilst also increasing flux to the lytic lysosome for degradation and an increase in catabolic supply to the cell. How eukaryotic cells respond to changes in nutritional levels can be controlled at different levels, but the mechanisms underlying many of these are not fully understood. Studies in yeast have revealed how large-scale surface protein degradation is mediated in response to restricted nutrients at late log phase (MacDonald et al., 2012) or relatively severe starvation conditions, such as lack of nitrogen or carbon (Lang et al., 2014; MacGurn et al., 2011; Müller et al., 2015). More specific nutrient starvation, such as depletion of vitamins, such as NAD^+^, or amino acids, such as Leucine, trigger different trafficking responses to increase ubiquitin-dependent trafficking of PM cargoes to the lysosome for degradation (Jones et al., 2012; MacDonald et al., 2015; MacDonald and Piper, 2017).

Cargo internalisation via clathrin mediated endocytosis is the best characterised mechanism to package cargo into endocytic vesicles and is governed by dozens of different factors. Although we have yet to fully understand how these factors cooperate, many aspects about the function (Kaksonen and Roux, 2018) and sequential coordination (Cocucci et al., 2012; Kaksonen et al., 2003; Mund et al., 2018) of these endocytic proteins has been revealed. The process is initiated by cytosolic adaptors, such as Assembly polypeptide 2 (AP2), which has been shown to coordinate cargo recruitment at specific lipid interactions sites, in addition to binding clathrin itself (Kelly et al., 2014). Other adaptors, such as epsin, clathrin assembly lymphoid myeloid leukemia protein (CALM) or its neuronal counterpart assembly protein 180 kDa (AP180), also interact with surface lipids through defined domains early in the endocytic process (Ford et al., 2001; Itoh et al., 2001; Miller et al., 2015). Recruitment of clathrin and assembly of these components alongside actin polymerisation and membrane bending Bin-Amphiphysin-Rvs (Bar) domain proteins serve to generate the burgeoning vesicle, followed by enzyme driven scission (Kaksonen and Roux, 2018). Additionally, mechanisms driving clathrin independent endocytosis have also been described (Mayor et al., 2014). Internalised proteins that retain a ubiquitination signal, largely mediated by the E3-ligase Rsp5 in yeast, are recognised by the Endosomal Sorting Complex Require for Transport (ESCRT) apparatus and delivered through the multivesicular body (MVB) pathway for degradation in the lysosome / yeast vacuole (Laidlaw and MacDonald, 2018). Proteins that are not destined for degradation can recycle back to the surface, including routes that return material to the surface via the *trans*-Golgi network (TGN) through dedicated machineries that interact with cargoes for recycling (McNally and Cullen, 2018). Recycling can also occur directly from early endosomes or indirectly, first traversing defined recycling endosomes (MacDonald and Piper, 2016). Endosomal organisation and recycling mechanisms in yeast are less clear (Ma and Burd, 2019), however, it was recently shown that although retrograde recycling of the yeast exocytic v-SNARE protein Snc1 is perturbed by deubiquitination (Xu et al., 2017), some cargo recycling is triggered by deubiquitination (MacDonald and Piper, 2017). Genetic dissection of this latter pathway implies recycling is controlled at the transcriptional and metabolic level, suggesting that early endocytic trafficking decisions contribute to the eventual downregulation of proteins during starvation.

Surface cargoes are also organised spatially within the plasma membrane. For example, in yeast, although membrane proteins are dispersed throughout the surface, certain cargoes, including nutrient transporters, also diffuse into PM invaginations termed eisosomes (Bianchi et al., 2018; Grossmann et al., 2008; Spira et al., 2012). Eisosomes have recently been shown to regulate both lipids and proteins at the surface in response to stress (Babst, 2019). Yeast cells lacking their cell wall rely on eisosomes as a lipid reservoir to allow PM expansion during hypoosmotic stress, as has been shown for mammalian caveolae (Kabeche et al., 2015; Sinha et al., 2011). Plasma membrane tension is sensed by the eisosomal osmotic stress sensors Slm1/2, which subsequently activate TORC2 to alter lipid metabolism (Riggi et al., 2018). Although, recent evidence on intact cells has suggested that although eisosomes might contribute to changes in lipids and membrane tension, there is no significant contribution of lipids to expanding cells during osmotic-stress (Appadurai et al., 2019). As to surface protein regulation, many nutrient transporters have been shown to localise to eisosomes, in particular during nutrient stress conditions when eisosomes maintain higher levels of these cargoes (Appadurai et al., 2019; Gournas et al., 2018; Grossmann et al., 2008; Moharir et al., 2018; Spira et al., 2012). Addition of transporter substrate results in a shift of transporters from eisosomes to other membrane compartments where they can function to uptake nutrients into the cell and subsequently be endocytosed and downregulated (Babst, 2019). It is not fully understood how eisosomes restrict access to endocytosis, but this capacity provides a protective compartment for cargoes to be preserved during mass downregulation.

Yeast has been a useful model to study physiological changes induced in response to changes in carbon source (Broach, 2012). Yeast preferentially uses glucose as a carbon source for fermentive growth, as with some rapidly growing mammalian cells (Diaz-Ruiz et al., 2011), but has also developed strategies to utilise various alternative carbon sources. During fermentive growth in replete glucose, alternative carbon utilisation pathways, for example galactose (*GAL*), maltose (*MAL*) and sucrose (*SUC*) genes, are actively repressed (Gancedo, 1998). The glucose sensitive transcriptional repressor Mig1, which binds to consensus sequences in the promoter regions of these example genes is responsible for this transcriptional repression (Griggs and Johnston, 1991; Hu et al., 1995; Nehlin and Ronne, 1990; Vallier and Carlson, 1994). Additionally, predicted Mig1 consensus binding sequences have been identified in an array of other functionally diverse genes (Wollman et al., 2017). Gene expression profiles in mutant cells lacking *MIG1* and/or the related repressor *MIG2* also span various functional classes beyond sugar metabolism (Westholm et al., 2008). When glucose starved cells are returned to glucose, alternative carbon transporters are endocytosed and rely on ubiquitination for degradation, which is provided by spatially distinct, cognate Arrestin-related Trafficking Adaptors (ARTs) at the surface and endolysosomal system (Becuwe et al., 2012; Becuwe and Léon, 2014; Hovsepian et al., 2018). Recent work has also shown how high affinity glucose transporters, which are not required during glucose starvation, are specifically endocytosed and degraded via Rsp5 and the Csr2 ART adaptor, which is repressed by Mig1/Mig2 in glucose rich conditions (Hovsepian et al., 2017). It is less clear if such transcriptional and post-translational regulatory mechanisms described for these sugar transporters during glucose starvation also control other surface membrane proteins. However, many distinct cargoes are downregulated when cells are shifted to media lacking a carbon source (Lang et al., 2014).

In this study, we reveal regulatory mechanisms of membrane trafficking pathways at the transcriptional level in response to glucose privation. We find a role for Mig1 mediated control of two distinct trafficking pathways: increasing internalization via clathrin mediated endocytosis and suppressing endocytic recycling to the surface. This control serves to degrade proteins more efficiently in response to acute depletion of glucose and provide the cell with energy to maintain viability during nutritional stress. *We* also find transcriptional control of eisosomes during the same glucose depleted conditions. Although not mediated through Mig1, eisosomal factors are transcriptionally upregulated and specific eisosomes increase in size to sequester a small portion of nutrient transporters. We propose a model where this reserve pool of surface localised nutrient transporters provide a physiological benefit upon a return to nutrient replete conditions, which outweighs the advantages of degrading an entire population of existing nutrient transporters.

## RESULTS

### Glucose depletion triggers cargo downregulation

As carbon source withdrawal inhibits translation initiation (Ashe et al., 2000), we provide the trisaccharide raffinose as an alternative sugar during glucose starvation experiments, to minimise additional stress whilst inhibiting glucose-dependent signal transduction pathways. The methionine permease Mup1, which is predominantly localised to the cell surface when cells are grown to logphase in media lacking methionine (MacDonald et al., 2012b), is a useful reporter for endocytosis. To monitor changes in Mup1-GFP localisation following replacement of glucose to raffinose media, we developed a microfluidic system that allowed rapid media exchange with simultaneous imaging. We found that extended incubations in glucose media, or repeated exchanges of fresh glucose media, had very little effect on the surface localisation of Mup1 (**Figure 1A**). However, upon glucose starvation Mup1-GFP is internalised and sorted through the endocytic pathway efficiently, accumulating inside the cell over time with almost no remaining signal at the plasma membrane after ~45 minutes. The trafficking kinetics of endocytosis upon glucose depletion are similar to substrate induced endocytosis of Mup1, triggered by addition of methionine (**Figure S1a**), suggesting glucose depletion serves to endocytose surface proteins at a rate close to the physiological maximum. Photobleaching during time-lapse microscopy did not affect endosomal GFP-tagged cargo but did reduce observable vacuolar signal in the confocal plane (**Figure S1b**). Therefore, Mup1 endocytosis and delivery to the vacuole for degradation was also assessed biochemically, with immunoblots of vacuolar processed GFP showing a large increase upon glucose starvation (**Figure 1B, 1C**). This raffinose exchange protocol also induced the downregulation of functionally and structurally distinct surface proteins, including the G-protein couple receptor Ste3 tagged with GFP, the uracil permease Fur4 tagged with mNeonGreen (mNG), and the arginine permease Can1 tagged with GFP (**Figure 1A, S1c**). However, we did note the ATP-binding cassette (ABC) transporter Yor1, which is downregulated in response to NAD^+^ starvation (MacDonald et al., 2015), showed no increase in endocytosis, even at extended raffinose incubations (**Figure S1d**). Furthermore, although relatively long exposures to lactate media triggers vacuolar delivery of high affinity hexose transporters tagged with GFP, we observed no sorting of Hxt6/7-GFP during raffinose treatment (**Figure S1e**). This suggests that glucose depletion simulates rapid downregulation of many, but not all, surface proteins. Raffinose treatment rapidly triggers relocalisation of Mup1-GFP to bright endosomal puncta at very short time points (e.g. 4 minutes), prior to sorting to the vacuolar lumen (**Figure 1D**). Additionally, endocytic uptake of FM4-64 at short time points was assessed in cells expressing a surface localised mutant of GFP-Snc1^PM^ (Lewis et al., 2000). Although raffinose treated cells display lower levels of FM4-64 dye binding (**Figure S1f**), possibly due to changes in the lipid composition of the plasma membrane induced upon glucose starvation, we find endocytosis to be efficient in glucose depleted conditions, with significant FM4-64 internalised following a 4 minutes uptake followed by quenching of extracellular dye (**Figure 1E**). We hypothesised this large-scale downregulation of surface proteins was controlled by a combination of increased endocytosis and decreased plasma membrane recycling.

**Figure 1:**
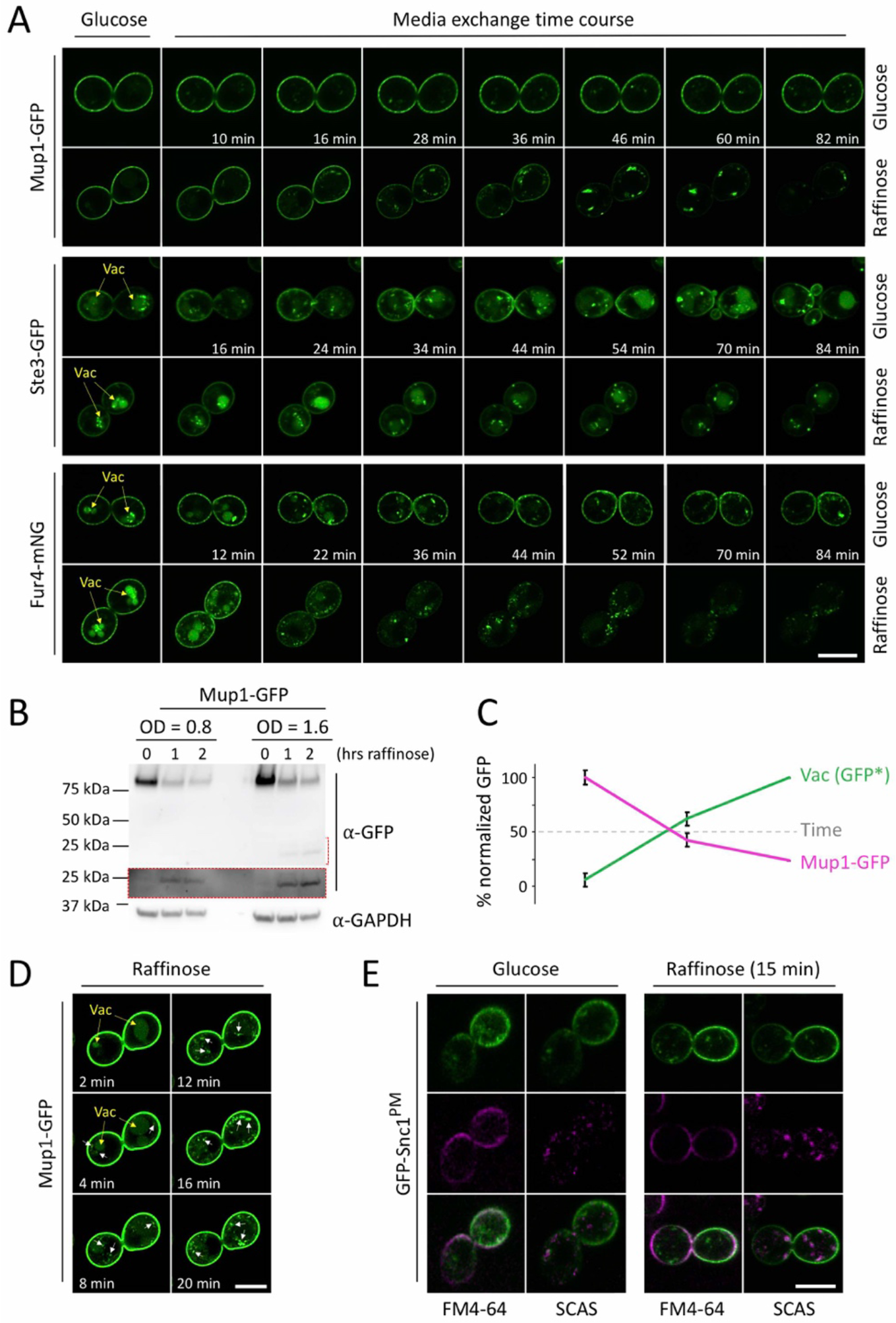
Glucose starvation results in downregulation of surface proteins: **A)** Wild-type cells expressing labelled cargoes (Mup1-GFP, Ste3-GFP, and Fur4-mNG) were grown to mid-log phase, processed for time-lapse microscopy and then imaged every 2-minutes in either glucose or raffinose media for indicated time course. Vacuoles are indicated with yellow arrows. **B)** Levels of Mup1-GFP in glucose and raffinose treated cells were assessed by immunoblotting lysates with GFP antibodies. Increased exposure (red box) shows levels of vacuolar processed GFP. Loading was assessed with anti-GAPDH antibodies. **C)** Graph depicts levels of Mup1-GFP and vacuolar processed GFP following raffinose addition at indicated times. Error bars show standard deviation from three replicate experiments. **D)** Increased exposure of early time points from experiment **(A)** to show internalised Mup1-GFP (white arrows). **E)** Wild-type cells expressing GFP-Snc1^PM^ were grown to mid-log phase in glucose media, adhered to a glass-bottom dish and incubated with glucose or raffinose for 30 minutes prior to flushing with FM4-64 media for 5 minutes, then exchanging with 2.4 μM SCAS media and imaging. Scale bar, 5 μm

### Mig1 regulates endocytic genes during starvation

We investigated a potential role for the Mig1 transcriptional regulator, which represses many genes in glucose replete conditions (Nehlin and Ronne, 1990; Nehlin et al., 1991), in cargo endocytosis following glucose starvation. Mig1 repression in the nucleus is alleviated by its translocation to the cytoplasm upon media exchange for alternative carbon sources (Vit et al., 1997). As expected, microfluidic exchange of media shows nuclear localisation is very sensitive to glucose availability, with rapid translocation to the cytoplasm following raffinose (**Figure 2A**) and return to nucleus when glucose is reintroduced (**Supplemental Video 1**). Mig1 binding sequences have been located to the promoter regions of approximately 100 yeast genes (Lundin et al., 1994; Wollman et al., 2017). In silico analysis of these candidates revealed several clusters, based on function or sub-cellular location, including many factors associated with protein downregulation (**Figure 2B**), including roles in ubiquitination, autophagy, and vacuolar degradation (**Supplemental Table 1**). Additionally, several endocytosis factors were identified as potential Mig1 targets (**Figure 2C**), including the genes encoding clathrin adaptors (*YAP1801, YAP1802*, and *APL3*) and the eisosome component PIL1, which is indirectly related to endocytosis (Babst, 2019; Goode et al., 2015). We also noticed that other endocytic genes contained consensus Mig1 binding sequences in their open-reading frame (ORF) sequences, so included these in downstream transcriptomic analyses.

**Figure 2:**
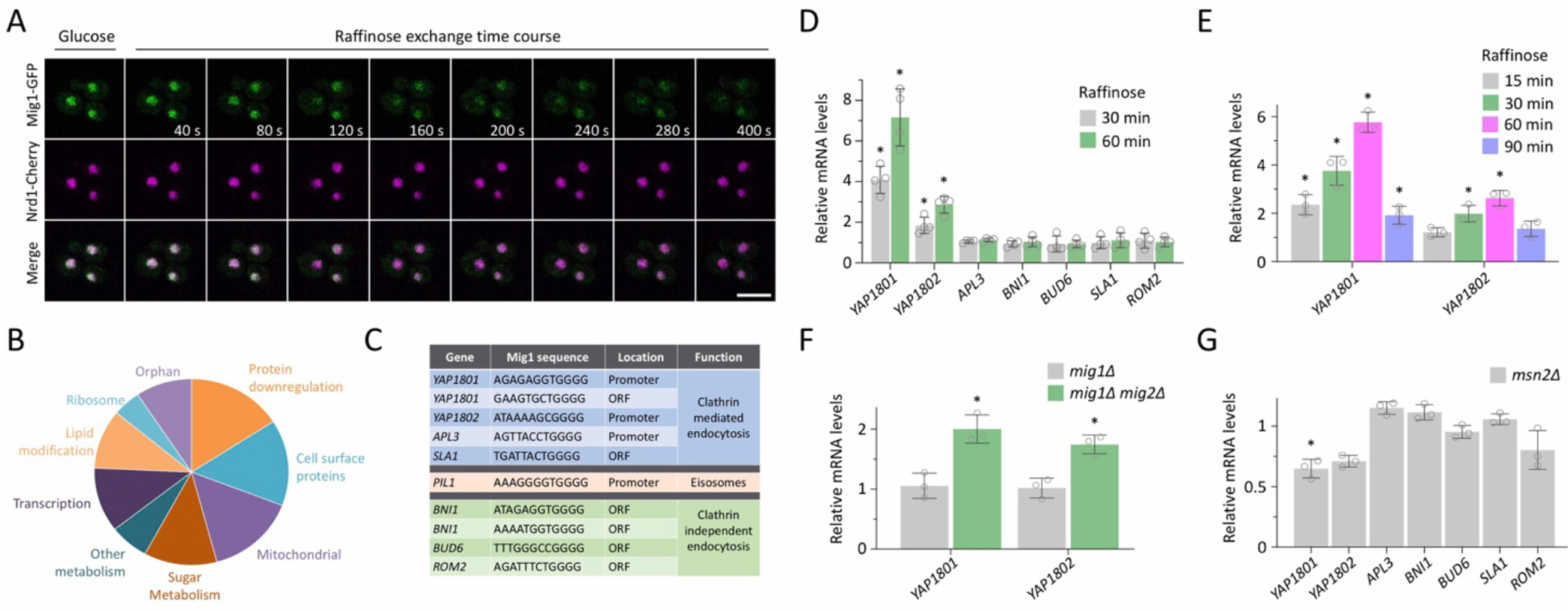
Mig1 and glucose regulates expression of clathrin adaptor genes: **A)** Time-lapse microscopy of wild-type cells co-expressing Mig1-mGFP and Nrd1-mCherry grown in glucose media prior to exchange with raffinose media for indicated times. **B)** Pie chart showing in silico categorisation of Mig1 binding targets in the yeast genome. **C)**Table of putative Mig1 target genes previously shown to function in endocytosis. **D - G)** Total RNA was prepared and transcript levels measured by quantitative RT-PCR of indicated genes, using ACT1 and HEM2 genes as a control, from indicated cells grown to mid-log phase in glucose media. Comparing raffinose exchange for 30 and 60 minutes **(D)** or 15, 30, 60 and 90 minutes **(E)** with glucose controls or comparing mig1 and mig1Δ mig2 cells **(F)** and msn2Δ cells with wild type **(G)**. Data shown is from 3 – 4 biological replicates (3 technical replicates) with error bars representing standard deviation. * indicates t-test p-values <0.05. Scale bar, 5 μm

To test if these potential Mig1 target genes were regulated in response to glucose availability, qPCR conditions were optimised with various primer pairs for each gene of interest (**Figure S2a - c**) and the testing of different housekeeping reference genes (Teste et al., 2009) during raffinose treatment. Most candidates tested were unchanged, however, we found genes encoding the yeast AP180 clathrin adaptors were upregulated in raffinose media. Most prominently, *YAP1801* exhibited a robust ~4-fold increase in expression levels 30 minutes after raffinose addition, which increased to ~6 fold at 60 minutes (**Figure 2D**). *YAP1802* levels also increased significantly, with a sustained ~2-fold increase following starvation incubations. Transcriptional upregulation of yeast AP180 adaptors was relatively acute, with transcript levels returning close to basal 90-minutes after raffinose (**Figure 2E**). However, as Yap1801 and Yap1802 are relatively stable, with halflife estimates of 3.6 hours and 6.1 hours, respectively (Christiano et al., 2014), we presume these transcriptional changes are sufficient to contribute to increased cargo endocytosis observed during nutrient stress (**Figure 1**). We find that *mig1*Δ *mig2*Δ cells show elevated levels of *YAP1801* and *YAP1802* transcript levels when compared to wild-type cells (**Figure 2F**). There was no appreciable change in clathrin adaptor levels in *mig1*Δ cells, with it being necessary to also delete *MIG2*, which has many of the same regulatory elements as Mig1 (Westholm et al., 2008) but is not translocated from the nucleus in responsive to glucose depletion (**Figure S2d**). In further support for a role of Mig1 in *YAP1801/02* expression, we also find decreased transcript levels in *msn2*Δ null cells, which lack the Msn2 regulator that is known to activate many of the genes that Mig1 represses (Lin et al., 2015).

### Glucose sensitive AP180 adaptors trigger endocytosis

The AP180 proteins are conserved early acting endocytic factors that bind phosphatidylinositol 4,5-bisphosphate at the plasma membrane, via their AP180 N-terminal homology (ANTH) domain, to recruit clathrin to endocytic sites (Ford et al., 2001; Wendland and Emr, 1998). We endogenously tagged the yeast AP180 adaptors with mGFP and confirmed that raffinose exchange results in a significant increase in Yap1801-mGFP proteins levels detected by immunoblot (**Figure 3A, B**). A modest change was observed for Yap1802-mGFP, which mirrors the modulation in transcript levels (**Figure 2**) and suggests *YAP1802* is less glucose sensitive. We find that endogenously tagged versions of AP180 adaptor proteins show punctate surface localisation, with both distinct and over-lapping localisations (**Figure 3C**). These confocal experiments indicated an increase in fluorescence intensity following acute raffinose treatment (**Figure S3b - d**) but we sought a more accurate method to study endocytosis in living yeast cells. For this, we employed single-molecule Slimfield (**Figure S3e**), using a narrow field of laser excitation (Wollman and Leake, 2016), which enabled precise spatial resolution and millisecond time sampling (Badrinarayanan et al., 2012; Plank et al., 2009; Reyes-Lamothe et al., 2010). Levels of endogenously tagged Yap1801 and Yap1802 were estimated to be ~1000 and ~400 molecules per cell, respectively, similar to previously documented estimates (Ho et al., 2018). There is a small but significant increase in abundance for both Yap1801 and Yap1802 following glucose starvation (**Figure 3D**). Endogenously mCherry labelled versions of Yap1801/02 were then used to monitor their localisation in relation to cargo in cells co-expressing Mup1-GFP. In replete conditions, Mup1 levels are dispersed across the plasma membrane, overlapping with spots of both Yap1801 and Yap1802 (**Figure 3E**). Most Mup1-GFP signal is internalised following raffinose treatment (**Figure 1**), however Airyscan confocal microscopy shows that although no longer dispersed across the surface, Mup1-GFP was still found in punctate structures decorated with Yap1801/02. Other fluorescent spots may also be explained by Mup1-GFP localisation in eisosomes, discussed later.

**Figure 3:**
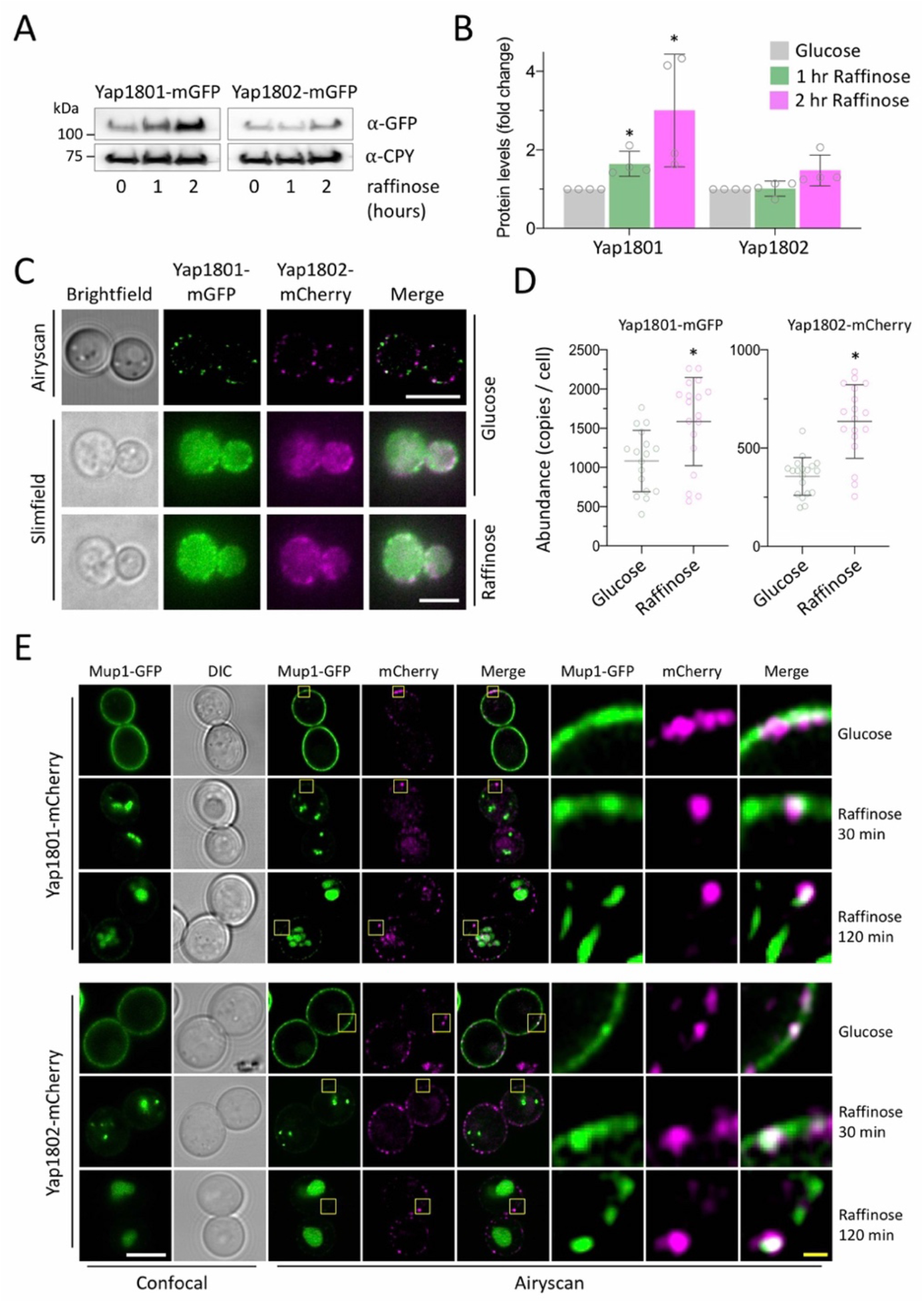
Yap1801 / 02 proteins are upregulated during glucose starvation: **A)** Immunoblot analysis of lysates generated from wild-type cells expressing endogenously tagged Yap1801-mGFP and Yap1802-mGFP in glucose and raffinose media. **B)** Histogram showing average Yap1801-mGFP and Yap1802-mGFP intensity compared to glucose over 3 biological replicates, with error bars showing standard deviation. **C)** Wild-type cells expressing Yap1801-mGFP and Yap1802-mCherry were grown to mid-log phase before processing for imaging by Airyscan confocal and Slimfield microscopy. **D)** Numbers of Yap1801 and Yap1802 molecules per cell were estimated using autofluorescence and background corrected integrated density values obtained via ImageJ software. **E)** Airyscan confocal co-localisation of Mup1-GFP with Yap1801-mCherry (upper) and Yap1802-mCherry (lower) in glucose and raffinose media. Yellow box denotes zoomed in regions. *indicates t-test p-values <0.05. White scale bar, 5 μm; yellow scale bar, 1 μm.

The model predicts that elevated levels of Yap1801 and Yap1802 induced upon glucose starvation are sufficient to upregulate endocytosis. To test this, mCherry tagged AP180 adaptors were expressed from a plasmid under the control of the copper inducible *CUP1* promoter. With no addition of copper, low levels of Yap1801/02-mCherry, similar to endogenously tagged versions, had no effect on the surface localisation of co-expressed Mup1-GFP (**Figure 4A**). However, over-expression of Yap1801-mCherry in glucose grown cells resulted in a significant portion of Mup1-GFP accumulating in endosomes. Over-expression of Yap1802-mCherry had a more profound effect, with many cells exhibiting Mup1-GFP localisation inside the lumen of the vacuole. Furthermore, Mup1-GFP endocytosis was induced slightly in *mig1*Δ cells, and substantially more in *mig1*Δ *mig2*Δ cells (**Figure 4B**), presumably through the increased levels of Yap1801/02. Similarly, the partial sorting of Mup1-GFP to the vacuole in cells grown to late log-phase (MacDonald et al., 2015, 2012) is also increased in *mig1*Δ *mig2*Δ mutants. Finally, we find an increase in the endocytosis of functionally distinct cargoes, including an increase in Can1-GFP sorting to the vacuole and a decrease in surface localised populations of both Ste3-GFP and Fur4-mNeonGreen, in *mig1*Δ *mig2*Δ cells (**Figure 4C**). Collectively, these data show that clathrin mediated endocytosis of cargo is triggered by Yap1801/02 in response to depleted glucose levels sensed by Mig1.

**Figure 4:**
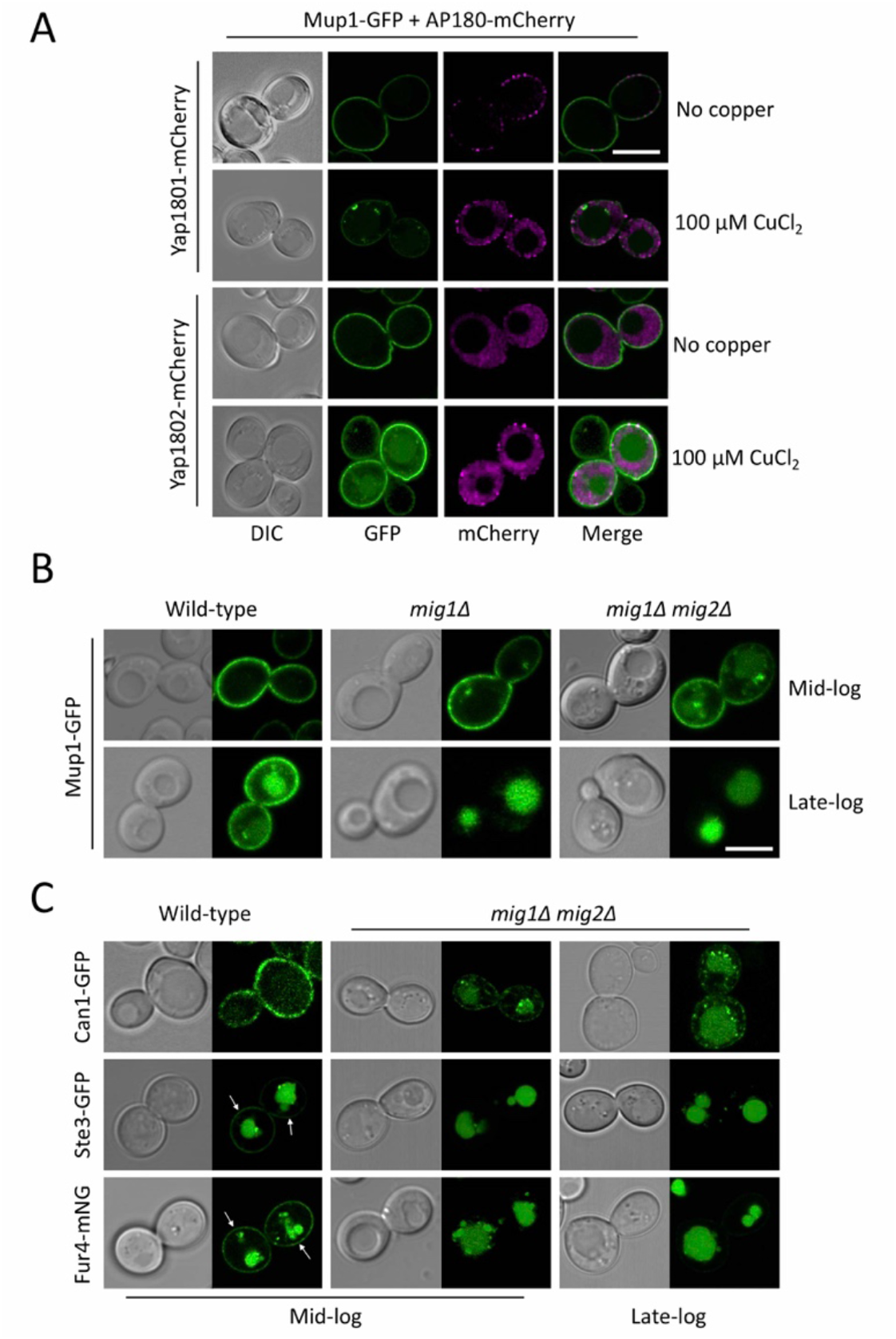
Increased Yap1801/02 increases rates of cargo endocytosis: **A)** Airyscan confocal microscopy imaging of wild-type cells coexpressing Mup1-GFP, from endogenous promoter, and Yap1801-mCherry or Yap1802-mCherry both expressed under control of the copper inducible CUP1 promoter using media lacking copper (upper) or upon addition of 100μM copper chloride for 2 hours. **B)** Localisation of Mup1-GFP expressed in wild-type, mig1Δ, and mig1Δ mig2Δ cells prepared for imaging at mid-log and late-log phase. **C)** Wild-type and mig1Δ mig2Δ cells expressing Can1-GFP, Ste3-GFP and Fur4-mNG were grown to mid-log and late-log phase before harvesting and preparation for fluorescence microscopy. Surface localised signal is indicated with white arrows. Scale bar, 5 μm.

### Galpha subunits control surface recycling in response to glucose

To assess if glucose starvation inhibited recycling of internalised material, to further promote vacuolar degradation, model protein and lipid recycling cargoes were analysed (**Figure 5A**). Efflux of internalised lipid dye FM4-64 can be used to measure the rate of recycling (Wiederkehr et al., 2000), which has previously been used to show carbon source removal inhibits dye efflux (Lang et al., 2014). Similarly, we find shifting cells to raffinose also results in robust inhibition of recycling (**Figure 5B**).

**Figure 5:**
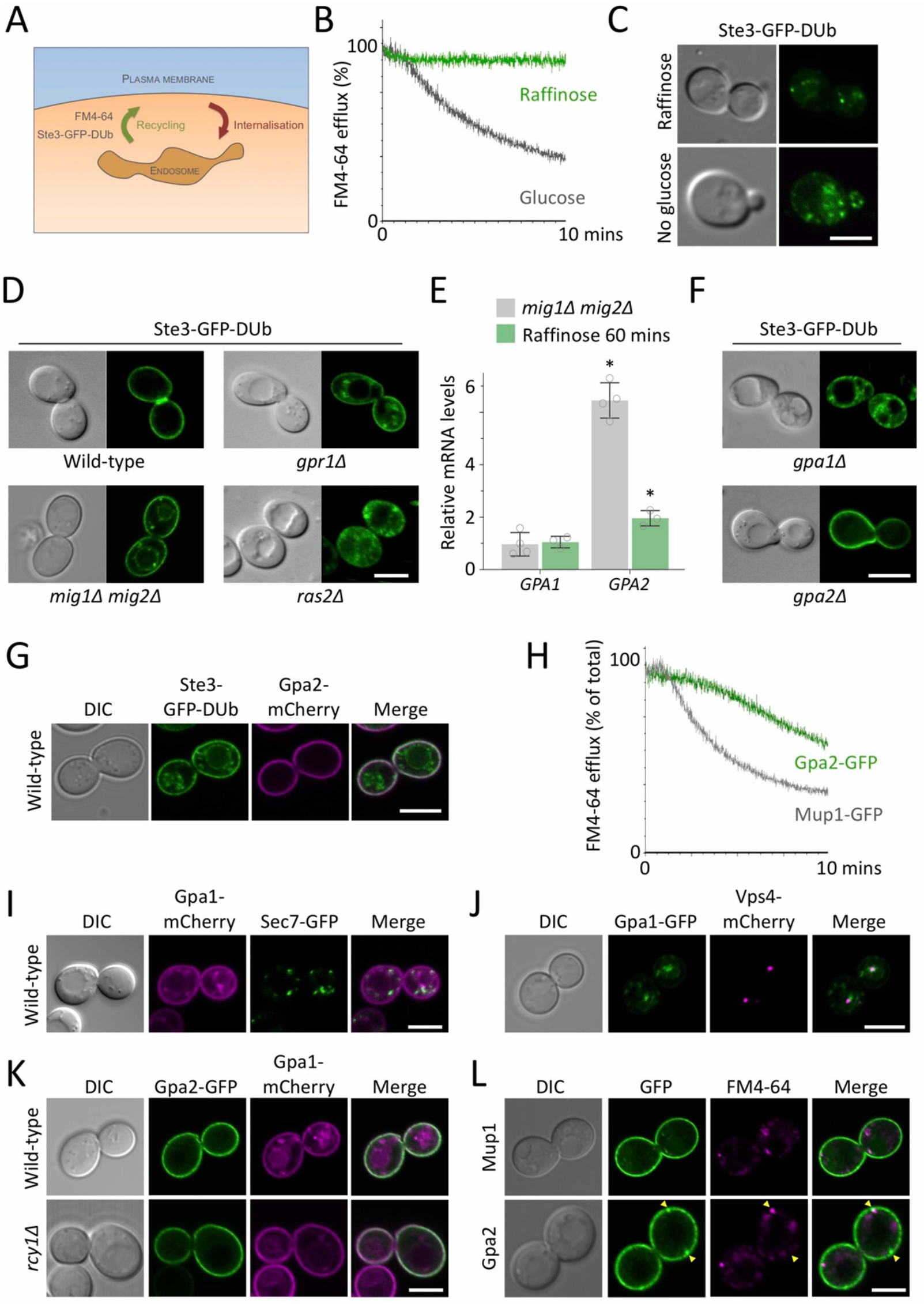
Glucose levels regulate endosomal recycling through Gpa1 and Gpa2: **A)** Schematic representation of reporter cargoes, the lipid dye FM4-64 and the engineered fusion of Ste3-GFP with a deubiquitinating enzyme (DUb), which are internalised and efficiently recycled. **B)** FM4-64 efflux measurements from wild-type cells grown in glucose or 30 minutes raffinose media prior to loading with dye for 8 minutes at room temperature. **C)** Ste3-GFP-DUb localisation was recorded in cells treated with media supplemented with raffinose (upper) or lacking carbon source (lower). **D)** Stably integrated Ste3-GFP-DUb was expressed in indicated strains, grown in SC media containing glucose and localisation confirmed by fluorescence microscopy. **E)** Relative mRNA levels of GPA1 and GPA2 transcripts, as measured by quantitative RT-PCR, are shown from 3 biological replicates in mig1Δ mig2 cells (grey) and following 60 minutes raffinose treatment (green). Error bars show standard deviation. **F)** Indicated mutants expressing Ste3-GFP-DUb were grown to log phase before preparation for confocal microscopy. **G)** Wild-type cells expressing Ste3-GFP-DUb were grown in 100 μM copper chloride to induce co-expression of Gpa2-mCherry under control of the CUP1 promoter. **H)** Wild-type cells expressing either Mup1-GFP (as a control surface protein) or Gpa2-GFP were loaded with FM4-64 for 8 minutes before washing and flow cytometry efflux measurements recorded. **I - K)** Wild-type cells expressing Gpa1-mCherry from the CUP1 promoter in the presence of 50 μM copper chloride and endogenously expressing Sec7-GFP **(I)**, Vps4-GFP **(J)** or co-transformed with a plasmid expressing Gpa2-GFP **(K)**. **L)** Cells expressing either Mup1-GFP or Gpa2-GFP were labelled with 40 μM FM4-64 for 3 minutes followed by quenching of extracellular dye with 2.4 μM SCAS and confocal imaging. * indicates t-test p-values <0.05. Scale bar, 5 μm.

We have previously developed a protein reporter for endocytic recycling, based on a modified version of the G-protein couple receptor (GPCR) Ste3 tagged with GFP and fused to the catalytic domain of a deubiquitinating enzyme (MacDonald and Piper, 2017; Stringer and Piper, 2011). Efficient recycling of Ste3-GFP-DUb is blocked in media either lacking sugar or supplemented with raffinose, which results in a rapid accumulation of the reporter in intracellular endosomal structures (**Figure 5C**). These results suggest a glucose sensitive signal transduction mechanism might also be at play. To test if glucose control of recycling could be traced to Mig1/Mig2 activity, Ste3-GFP-DUb was expressed in *mig1*Δ *mig2*Δ mutants to reveal recycling is defective in these cells (**Figure 5D**). These accumulations might be explained by increased internalization via Yap1801/02, but as the elevated internalization of Ste3 upon addition of **a**-factor does not accumulate Ste3-GFP-DUb in endosomes (MacDonald and Piper, 2017), we propose *mig1*Δ *mig2*Δ cells are also defective in recycling.

To identify potential machinery that controls recycling via Mig1 in response to glucose depletion, we compared putative Mig1 target genes (**Supplemental Table 1**) with defective recycling mutants identified from a genetic screen (MacDonald and Piper, 2017). Simply cross-referencing these lists did not yield any matches, as might be expected as we sought a Mig1-repressed factor that *inhibits* recycling upon *increased* expression. We next surveyed the 89 recycling mutants for associations with glucose metabolism and found glucose sensing GTPases Gpr1 and Ras2 are required for recycling, which we confirmed with stable integration of Ste3-GFP-DUb in the respective null strains (**Figure 5D**). Intriguingly, *GPA2*, which encodes the Galpha subunit Gpa2 that functions with Gpr1 to induce cAMP signal upstream of Ras (Kraakman et al., 1999; Xue et al., 1998), was identified as a candidate gene repressed by Mig1. Furthermore, although the other yeast Galpha subunit Gpa1, which functions in the pheromone response pathway but is not linked to Gpr1 activity (Dietzel and Kurjan, 1987; Miyajima et al., 1987), it is required for recycling (MacDonald and Piper, 2017). Levels of *GPA1* are not regulated by glucose or Mig1 but *GPA2* is transcriptionally upregulated ~2-fold in *mig1*Δ *mig2*Δ cells and ~6-fold upon a shift to raffinose for 1 hour (**Figure 5E**). As over-expression of Gpa2 does not inhibit Gpr1 function, instead triggering increased Ras signalling and accumulation of cAMP (Nakafuku et al., 1988), we sought an alternative mechanism. Gpa2 physically and genetically interacts with Gpa1 (Ho et al., 2002; Xue et al., 1998), suggesting that increased levels of Gpa2 might interfere with the ability of Gpa1 to regulate recycling, supported by the fact that Ste3-GFP-DUb recycling is very defective in *gpa1*Δ cells but is unaffected in *gpa2*Δ mutants (**Figure 5F**).

These data support a model whereby Gpa1 plays an essential role in recycling, which is suppressed by elevated levels of Gpa2 during starvation. In support of this, over-expressing Gpa2-mCherry inhibits surface recycling of Ste3-GFP-DUb (**Figure 5G**). The role of Gpa1 in recycling is not specific to the pheromone pathway GPCR Ste3 based reporter, as over-expressing Gpa2-GFP also inhibits general recycling of lipids, measured by FM4-64 efflux (**Figure 5H**). As Gpa1 has been shown to associate with the yeast phosphatidylinositol 3-kinase (Vps34 and Vps15) and stimulate production of phosphatidylinositol 3-phosphate (Slessareva et al., 2006) we propose this lipid regulation is required for efficient recycling during replete glucose conditions. Indeed, Gpa1-mCherry localises to endosomal compartments marked by Ste3-GFP-DUb, which excessively accumulate in recycling deficient *rcy1*Δ mutants (**Figure S5a**). At steady state, some Gpa1-mCherry co-localises with late Golgi compartments marked with Sec7-GFP, whilst another population is distinct (**Figure 5I**). Previous studies have shown that a constitutively active Gpa1^Q323L^ mutant almost exclusively localises to late endosomes marked with ESCRTs (Slessareva et al., 2006), suggesting it may be inactive during residency at the Golgi and that the primary site of recycling is at late endosomal compartments, marked with the ESCRT associated factor Vps4 (**Figure 5J**). As mentioned, Gpa1 and Gpa2 physically interact (Ho et al., 2002), and we find significant colocalization of Gpa1-mCherry and Gpa2-GFP at the plasma membrane. Endosomal localisation of Gpa1-mCherry is reduced in recycling deficient *rcy1*Δ cells, with most signal colocalising with Gpa2-GFP at the PM, potentially showing the recycling role of Gpa1 functions downstream of Rcy1 (**Figure 5K**). Alternatively, as Gpa1 is both palymitoylated and myristoylated (Song et al., 1996; Song and Dohlman, 1996), its ability to regulate endosomal lipids with Vps34/Vps15 may be required for its correct localisation. Surface localisation of Gpa1 was unperturbed by its overexpression or deletion of *rcy1*Δ. However, we did observe accumulations of Gpa2-GFP at the PM marked by sites of FM4-64 trafficking, in a pattern not observed for surface proteins like Mup1-GFP (**Figure 5L**), and in transient intracellular compartments during time-lapse microscope (**Figure S5b**). These latter localisations failed to co-localise with Gpa1-mCherry or Sec7-GFP Golgi compartments, but did overlap with Vps4 endosomes (**Figure S5c**), suggesting Gpa2 play a more direct role in recycling in addition to sequestering Gpa1 at the PM

Collectively, we therefore conclude that the downregulation of cell surface membrane proteins in response to glucose starvation is achieved by combining an increase of endocytosis, mediated through higher levels of the yeast AP180 clathrin adaptors, with an increase in Gpa2, which serves to interfere with Gpa1-mediated lipid regulation required for recycling.

### Eisosomal reorganisation sequesters cargo during glucose starvation

Spatial mapping and functional analyses of surface cargoes have recently shown lipid domains, termed eisosomes, harbour surface proteins and antagonize their endocytosis (Bianchi et al., 2018; Busto et al., 2018; Gournas et al., 2018; Moharir et al., 2018). Eisosomal regulation of surface proteins in response to glucose starvation is implied from *PIL1*, which encodes the structural Pil1 protein required for the proper formation of eisosomes (Walther et al., 2006), being identified as a potential Mig1 target (**Supplemental Table 1**). As expected, fluorescently tagged Pil1 fails to co-localise with sites of endocytosis marked by Yap1801/02 (**Figure 6A**). Upregulation of eisosomes that contravene cargo endocytosis may seem contradictory in conditions that also upregulate endocytosis, however, cargo protection in eisosomes during starvation and osmotic shock has recently been documented (Appadurai et al., 2019; Gournas et al., 2018). To test if eisosomal factors are regulated at the transcriptional level, we tested expression levels of many genes associated with eisosomes in glucose versus raffinose media. The levels of *PIL1* and *PKH2* were increased by ~2-fold, but others showed more dramatic increases, with mRNA levels of *NCE102* and *LSP1* increasing ~6-fold and *SLM1* levels ~11-fold after 60 minutes treatment with raffinose (**Figure 6B**). We note that *mig1*Δ *mig2*Δ cells show no increase in *PIL1* levels, however, as most of the genes encoding eisosomal factors upregulated in raffinose are not predicted to be repressed by Mig1, additional transcriptional regulators must integrate with this nutrient stress response.

**Figure 6:**
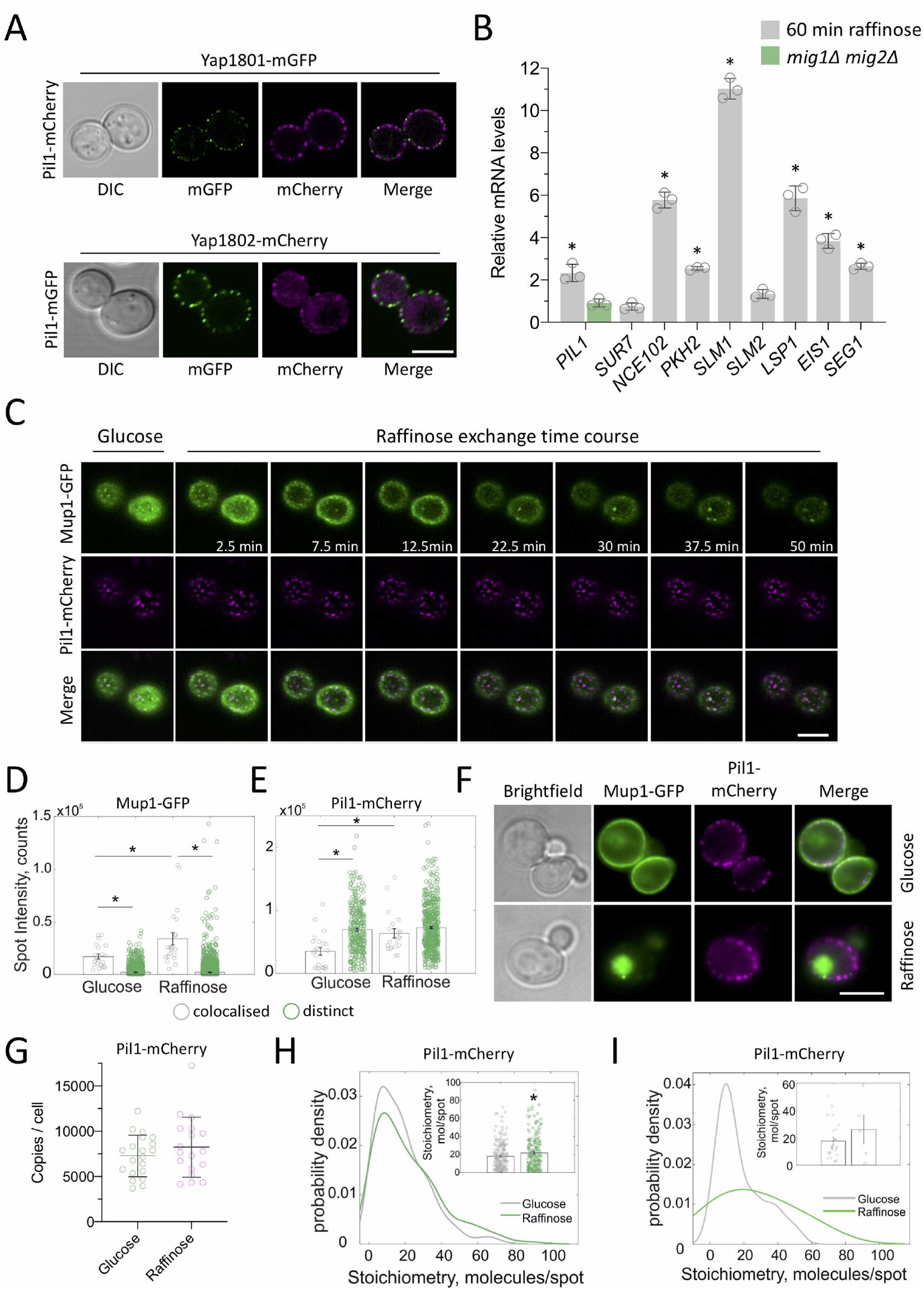
Eisosomes sequester surface proteins in glucose starvation conditions: **A)** Wild-type cells endogenously expressing Yap1801-mGFP transformed with a plasmid expressing Pil1-mCherry (left) or expressing a chromosomally tagged Yap1802-mCherry and Pil1-GFP from a plasmid were grown to mid-log phase and prepared for fluorescence microscopy. **B)** Quantitative RT-PCR of indicated genes, was performed from RNA extracted from wild-type cells grown in glucose media and relative levels compared to cells grown in raffinose media for 1 hour. Error bars show the standard deviation from 3 biological replicates (each averaged from 3 technical replicates). **C)** Average intensity projection of top focussed Airyscan confocal images shown for raffinose exchange time course of wild-type cells co-expressing Pil1-mCherry and Mup1-GFP. **D - E)** Jitter plots of fluorescence intensities of Mup1-GFP **(D)** and Pil1-mCherry **(E)** spots measured for both colocalised (grey) and distinct (green) foci from cells grown to mid-log phase and either prepared directly for confocal microscopy or first grown in raffinose media for 30 minutes. Error bars indicate standard error. **F)** Examples of Slimfield images of cells co-expressing Mup1-GFP and Pil1-mCherry in glucose (upper panel) and raffinose solid media conditions (lower panel). **G)** Number of Pil1-mCherry marked eisosomes per cell in glucose (n = 33 cells) and 30 minutes raffinose (n = 71 cells) and depicted with standard deviation error bars. **H)** Kernel density plot of Pil1-mCherry stoichiometry distribution in glucose (grey) and raffinose (green) conditions. Insert: jitter plot of Pil1-mCherry foci stoichiometry in glucose (grey) and raffinose (green) indicating means and standard error bars. **I)** Kernel density plot of stoichiometry distribution of Pil1-mCherry spots colocalised with Mup1-GFP in glucose (grey) and raffinose (green) conditions. Insert: jitter plot of Pil1-mCherry foci stoichiometry in glucose and raffinose indicating means and standard error bars. * indicates t-test p-values <0.05. Scale bar, 5 μm.

We assume this transcriptional response is relatively acute, as *PIL1* transcript levels peak around 60 minutes and return close to basal by 90 minutes (**Figure S6a**), although Pil1 is stable with half-life >6 hours (Christiano et al., 2014). To test if this transcriptional upregulation of eisosomes impacted cargo endocytosis we performed time-lapse microscope of cells co-expressing Mup1-GFP cargo and Pil1-mCherry, to mark eisosomes. We optimised 4D confocal conditions to best observe eisosomes at the surface (**Figure S6b**) before initiating glucose starvation by flushing raffinose media via microfluidics. Mup1-GFP co-localises with eisosomes in glucose replete media (**Figure 6C**), with the majority of Mup1-GFP being distributed at the plasma membrane. As shown, most Mup1-GFP is endocytosed following glucose starvation (**Figure 1**), however, we noticed concentration of Mup1-GFP within some, but not all, eisosomal structures (**Figure 6C**). This is in stark contrast to when the Mup1 substrate methionine is added, which triggers flux from eisosomes to distinct PM regions that allow internalisation (Busto et al., 2018), implying this response is related to starvation of the cell and not a mechanism associated with typical transporter regulation. It is curious that only certain eisosomes appear to accumulate Mup1-GFP, with no signal observed for other Pil1-marked compartments, suggesting a precise regulatory mechanism. In support of this idea, we captured dramatic reconfigurations of Pil1-mCherry in eisosomes retaining Mup1-GFP (**Supplemental Video 2**). To complement 4D confocal, a series of steady state experiments were performed to fully quantify this phenotype, which confirmed intensity of Mup1-GFP foci colocalised with eisosomes increased significantly following glucose starvation, with no change in intensity of Mup1-GFP molecules that were in other, non-eisosomal regions of the cell (**Figure 6D, S6c**). Similarly, the intensity of Pil1-mCherry was relatively unchanged following raffinose in regions that lack Mup1-GFP, but intensity increased more in eisosomes harbouring cargo (**Figure 6E**). We saw no change in eisosome number in cells shifted to raffinose media (**Figure S6d**).

To better understand these observations, we performed Slimfield microscopy of cells expressing Mup1-GFP and Pil1-mCherry during growth in both glucose and raffinose (**Figure 6F**). Using single particle tracking (**Figure S3e**), and previously optimised step-wise photobleach protocols (Leake et al., 2006; Shashkova and Leake, 2017) from micrographs acquired every 5 milliseconds enabling a lateral localization precision of ~40nm (Ford et al., 2001; Itoh et al., 2001; S. E. Miller et al., 2015). This analysis showed abundance of Pil1-mCherry was similar in each sugar (**Figure 6G**). Single-molecule analysis of plasma membrane localised proteins showed that although there were no physiological changes in Mup1-GFP (**Figure S6e, S6f**) the stoichiometry of Pil1-mCherry increased (**Figure 6H**), collectively suggesting that in response to starvation, eisososmes increase their size to better sequester cargo. Also, although most Mup1-GFP molecules are no longer found within the surface region following starvation, similar to confocal analysis, the few spots retained show that the stoichiometry of Pil1 in these regions increases (**Figure 6I**). Finally, we also noted a small but significant increase in diffusion coefficient estimates of eisosomal localised Pil1 molecules (**Figure S6g**). This might not be predicted for recruitment of soluble Pil1 molecules to a static eisosomal structures, however, may be explained by the molecular rearrangements of Pil1-mCherry we observed of these cargo-storing eisosomes (**Supplemental Video 2**). Collectively, this helps rationalise how a relatively modest transcriptional increase in Pil1, alongside more dramatic increases in other eisosomal factors, can be directed to a subset of eisosomes that serve to sequester Mup1-GFP during glucose starvation.

### Eisosomes provide a recovery growth benefit following starvation

Not all surface cargoes localise to eisosome compartments, with Pma1, for example, marking a distinct compartment region of the surface (Malínská et al., 2003). Even Ste3-GFP, which is robustly downregulated in response to raffinose (**Figure 1**), or Ste3-GFP-DUb that is almost exclusively localised to the plasma membrane showed no sign of colocalization with Pil1-mCherry marked eisosomes (**Figure S7a**). Our model to explain all these findings is that eisosomes increase in size specifically to better harbour nutrient transporters, to persist throughout the starvation period and provide a physiological benefit to the cell upon a return to replete conditions, allowing nutrient uptake without new transporter synthesis. We reasoned that cells deficient in eisosome function would exhibit growth deficiencies by virtue of their inability to maximise nutrient uptake. However, growth on rich and synthetic defined media and estimated doubling times, showed no obvious difference between wild-type and null cells lacking various eisosome components (**Figure 7A, S7b**). However, some mutants, including *pil1*Δ, *lsp1*Δ and *nce102*Δ fail to reach the optical density of wild-type cells upon growth to saturation (**Figure S7c**), implying their nutrient uptake capacity might be compromised. To more specifically test our model, we took advantage of the eisosomal localised protein Fur4 (Moharir et al., 2018), which when tagged with mNG shows a steady state localised population in the vacuole but 3D confocal also clearly shows retention within eisosomes (**Figure 7B**). Fur4 is a uracil transporter that is required for the uptake of uracil and analogues (Jund and Lacroute, 1970), which is required for growth of *ura3*Δ auxotroph strains in minimal media with low uracil concentrations (**Figure 7C**). We therefore reasoned that the ability of Fur4 to scavenge uracil following glucose starvation might provide a recovery-based assay with a dynamic range to test the role of eisosomes. For this assay, we used *pkh2*Δ cells, which reduce the levels of unmodified and phosphorylated Pil1 (**Figure 7D**) and induces Pil1 redistribution into reticular structures at the plasma membrane (**Figure 7E**) reminiscent of structures observed in *pkh2*Δ cells carrying a temperature sensitive allele of *pkh1* (Walther et al., 2007). We also used *eis1*Δ and *nce102*Δ cells that are defective in eisosome assembly (Aguilar et al., 2010; Fröhlich et al., 2009), which we also find disrupt Pil1 localisation. All of these mutants decreased the punctate localisation of Fur4-mNG to eisosomes marked by Pil1-mCherry (**Figure 7F**), allowing us to test the hypothesis that cargo storage in eisosomes aids recovery from nutrient stress. Cells were grown to early log phase in minimal media, before treatment with raffinose for two hours to trigger downregulation of surface proteins and accumulation of a subset of nutrient transporters in eisosomes. Cells were then returned to nutrient rich media of varying concentrations of uracil and growth recorded over time. The recovery rate of growth for wild-type cells was significantly better than all three null mutants with defective eisosomes, across all concentrations of uracil tested (**Figure 7G, S7d**). This demonstrates that the capacity of cells to position a portion of nutrient transporters in eisosomes during starvation conditions maximises nutrient uptake following a return to replete conditions.

**Figure 7:**
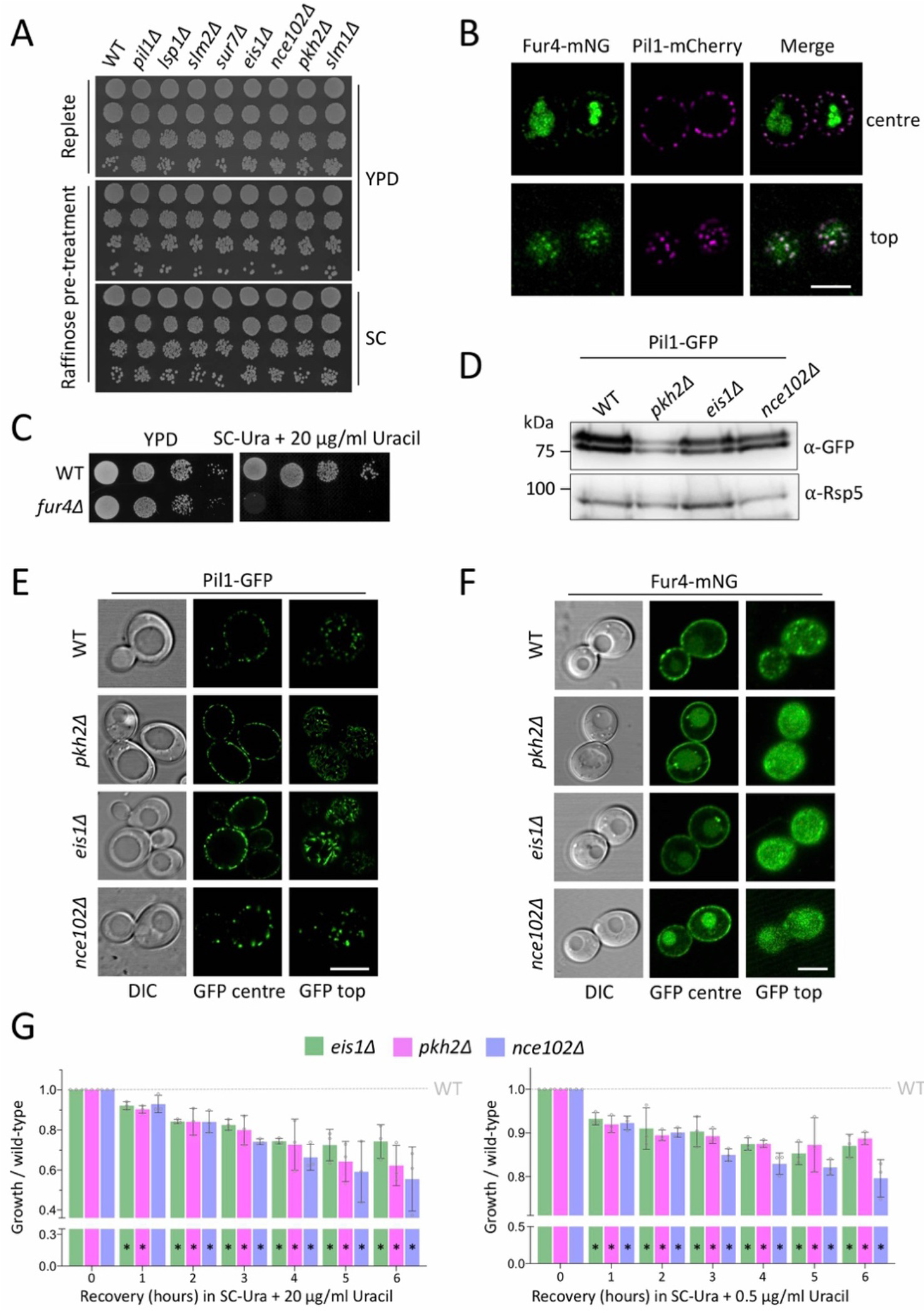
Eisosomes are required for efficient recovery following glucose starvation: **A)** Indicated yeast strains were grown to mid-log phase in glucose media and either prepared directly, or first incubated in raffinose media for 2 hours, before plating on YPD and SC media and growth recorded after two days at 30°C. **B)** Co-localisation of Pil1-mCherry and Fur4-mNGP by Airyscan confocal microscopy from cells grown to mid-log phase before imaging in minimal media focussed at the centre and top. **C)** Growth of wild-type and fur4Δ cells grown in rich media prior to serial dilution and growth on YPD and SC solid media for 2 days at 30°C. **D)** Levels of Pil1-GFP expressed from a plasmid in indicated strains were assessed by immunoblot of lysates generated from mid-log phase cells using anti-GFP antibodies and anti-Rsp5 antibodies as a loading control. **E)** Pil1-GFP localisation of transformants from **(D)** grown to log phase was recorded by confocal microscopy. **F)** Indicated strains were transformed with a plasmid expressing Fur4-mNG from its endogenous promoter and imaged by confocal microscopy. **G)** Indicated mutants were grown to mid-log phase in SC-complete media before harvesting and growth in raffinose media for two hours before recovery growth was assessed by OD_600_ measurements from cells incubated in SC media containing Uracil at 20 μg/ml (left) and 0.5 μg/ml (right). Error bars show standard deviation from 3 biological replicates. * indicates t-test p-values <0.05. Scale bar, 5 μm.

## DISCUSSION

Yeast cells provided with appropriate carbon and ammonium can synthesize all amino acids for protein translation, however, amino acid biosynthesis and transporter mediated uptake is coordinated with multiple metabolic programmes, including sugar utilization (Ljungdahl and Daignan-Fornier, 2012). We find the replacement of glucose with raffinose, a suboptimal alternative carbon source, induces the endocytosis and degradation of various surface proteins unrelated to sugar metabolism (e.g. amino acid transporters). We propose a glucose sensitive transcriptional response governs the activity of two endosomal membrane trafficking pathways, which act in concert to mediate cargo downregulation, including various nutrient transporters (**Figure 8**). Firstly, in glucose replete conditions endocytosis is maintained at a basal level, in part through the transcriptional repression via Mig1 (and Mig2) of the clathrin adaptor genes *YAP1801* and *YAP1802*. When glucose is exchanged for raffinose, Mig1 translocates from the nucleus resulting in elevated levels of cellular Yap1801/02, which is sufficient to accelerate endocytosis (**Figure 4**). We propose that increased yeast AP180 adaptors triggers a cascade of events, likely involving much of the canonical endocytic machinery, that work collaboratively to package and internalise an array of surface proteins. In addition to lipid binding domains, Yap1801 and Yap1802 associate with cortical clathrin (Newpher et al., 2005) and Pan1, which is required for endocytosis (Maldonado-Báez et al., 2008) and in turn activates Arp2/3 mediated actin assembly (Duncan et al., 2001; Wendland et al., 1999; Wendland and Emr, 1998). As the disordered domains of AP180 have been shown to amplify membrane curvature (Zeno et al., 2018), higher levels of these proteins are reasonable candidates for initiating a relatively broad downregulation of cargo following glucose replacement. We think this scenario best explains our observations of various cargoes being downregulated following increased levels of AP180 adaptors (in raffinose media, *mig1*Δ *mig2*Δ cells, and during plasmid borne over-expression), as deletion of these adaptors, as with many other yeast endocytic factors, does not have a strong defect in endocytosis (Huang et al., 1999; Wendland and Emr, 1998) or only a cargo specific defect for the internalisation of the SNARE protein Snc1 (Burston et al., 2009). Alternatively, increased sorting of Snc1, which is not a passive cargo but represents part of the core endocytic fusion machinery, may catalyse more general internalisation of other cargoes that are not necessarily interactors of Yap1801/02. We note that the increases in AP180 adaptor transcript levels in raffinose is relatively modest (between 1 and 6 fold), which presumably can coordinate meaningful upregulation of clathrin mediated endocytosis. This is in contrast to overexpression of AP180 (Ford et al., 2001) and its homologue CALM (Tebar et al., 1999) from plasmid borne CMV promoter constructs in mammalian cells, which inhibits endocytosis; we presume this mass over-expression serves to sequester clathrin in a manner disruptive to multiple processes, including internalisation. The finding that FM4-64 internalises efficiently in raffinose (**Figure 1E**), but not in sugar free media (Lang et al., 2014) may be explained by 1) lack of carbon source triggers shut down of internalisation to protect ATP maintenance and this is not necessary in the presence of raffinose, which can be hydrolysed by invertase or 2) changes in plasma membrane tension / osmotic pressure in media lacking sugar triggers TORC2 mediated changes in lipid composition (Appadurai et al., 2019; Riggi et al., 2018) that affect FM4-64 binding.

**Figure 8:**
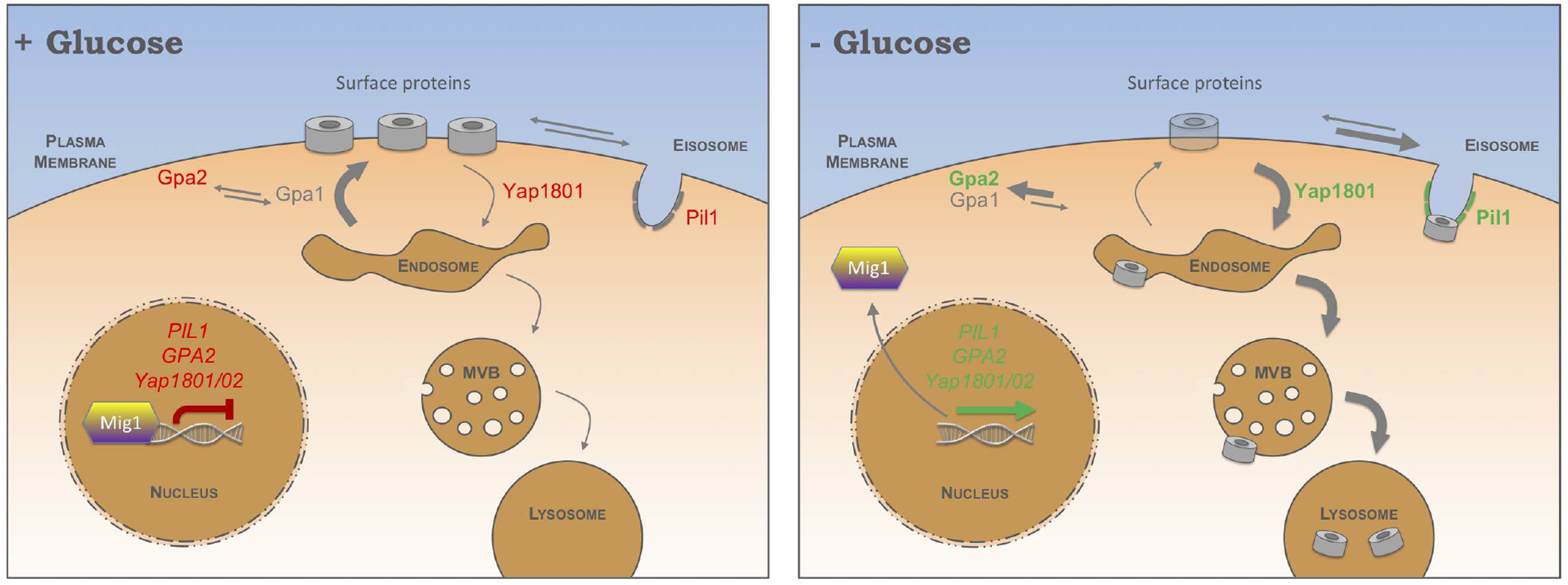
Model for modes of surface protein regulation in response to glucose: Schematic diagram shows the regulation of cell surface membrane protein cargoes in response to glucose replete (**left**) and glucose starvation (**right**) conditions. When cells have access to glucose for maximal growth, internalisation occurs at basal level with Mig1 repressing expression of adaptor genes YAP1801/02. In these conditions, recycling of internalised proteins efficiently returns many surface cargoes to the plasma membrane. This is in part through the Mig1 repression of GPA2 and the activity of Gpa1 localised to endosomes. In combination, this maintains high levels of cargo at the surface. Whilst resident at the surface, nutrient transporters diffuse in and out of eisosomes. Following a shift to raffinose, several metabolic changes occur. Firstly, Mig1 translocates from the nucleus, resulting in increased levels of Yap1801/02, which trigger increased cargo endocytosis. In raffinose conditions, Mig1 de-repression of GPA2 serves to inhibit recycling of protein and lipid cargo back to the surface. Modulation of these endosomal trafficking routes increases retention of cargo in endosomes and drives their ubiquitin-mediated trafficking to the lysosome for degradation. Although most nutrient transporter pools are degraded through the above described mechanisms, a small reserve population is sequestered in eisosomes during nutrient stress, which can be deployed upon a return to glucose replete conditions for efficient recovery following the period of starvation.

Under glucose replete conditions, clathrin mediated endocytosis of surface cargoes is counterbalanced by efficient recycling back to the surface, in combination maintaining high levels of surface proteins at the plasma membrane for optimal growth (**Figure 8**). Conceptually, a decrease in recycling to the surface would complement increased internalisation to maximise cargo sorting to the MVB pathway for degradation. We reveal a novel role Mig1/Mig2 in surface recycling in glucose conditions, which repress the expression levels of *GPA2* (**Figure 5E**). Increased levels of Gpa2, such as those found in raffinose grown cells, is sufficient to reduce recycling efficiency. To rationalise these observations at a mechanistic level, we confirmed that deletion of several glucose sensing factors, identified in a previous screen (MacDonald and Piper, 2017), results in defective recycling. The role of Gpa1 in protein and lipid recycling could be explained by a model presented by Dohlman and colleagues (Slessareva et al., 2006), whereby Gpa1 has a role outside of surface GPCR signalling and can functionally connect with the yeast phosphatidylinositol 3-kinase (Vps34/Vps15). As Vps34 / Vps15 activity produces phosphatidylinositol 3-phosphate (PtdIns 3-P), which is required for various post-Golgi membrane trafficking events (Reidick et al., 2017), we propose lipid regulation via Gpa1 attenuates recycling in response to glucose starvation. The finding that Gpa1 physically binds Gpa2 (Ho et al., 2002), suggests these observations could be functionally connected. In glucose media, relatively lower levels of Gpa2 at the plasma membrane maintained via Mig1 repression, would allow Gpa1 to remain unfettered in its role at endosomes recycling membrane proteins back to the surface. Vps34 / Vps15 were identified as Gpa1 signalling partners from a systematic genome-wide screen for deletion mutants that attenuate the negative effects of constitutively active Gpa1 (Slessareva et al., 2006); the top hit from this screen was *reg1*Δ mutants. Reg1 is a phosphatase that works upstream of Mig1, therefore *reg1*Δ cells phenocopy *mig1*Δ *mig2*Δ mutants (Schüller, 2003). These observations support our model that Gpa1-Gpa2 regulates recycling from endosomes back to the surface via Vps34/Vps15 signalling. A further connection was revealed as recycling defective *rcy1*Δ mutants mislocalise Gpa1 (**Figure 5K**), suggesting Gpa1-Vps34-Vps15 signalling is disrupted or fails to initiate in *rcy1*Δ cells, which do not recycle material back to the surface efficiently (Galan et al., 2001). Although we have not defined the mechanism by which *gpr1*Δ and *ras2*Δ mutants attenuate recycling of Ste3-GFP-DUb, it may be that they are related to the balance of action between Gpa1-Gpa2. Alternatively, these factors might work in a distinct pathway, indeed it has been shown that protein kinase A (PKA) alongside Gpr1 and Ras signalling is related to alterations in post-Golgi trafficking factors in response to glucose starvation (Aoh et al., 2011).

The finding that elevated endocytosis and inhibition of recycling work together to downregulate surface proteins conceptually aligns with a starvation model to degrade existing resources to supply essential processes whilst turning off non-essential processes. Modulating both internalisation and recycling pathways to increase endosomal retention of surface proteins will enhance cargo ubiquitination at late endosomes, which is sufficient to trigger ESCRT recruitment, luminal vesicle formation and subsequent cargo sorting through the MVB pathway (MacDonald et al., 2012b). Endosomal cargo ubiquitination is largely mediated by the E3-ligase Rsp5 in complex with cargo-specific adaptors, some of which are also regulated in response to glucose via adenosine monophosphate-activated protein kinase (AMPK) (O’Donnell and Schmidt, 2019). We have recently shown that Rsp5 adaptors compete against one another for occupancy on Rsp5 to direct activity to their cognate substrates(MacDonald et al., 2020). As certain Rsp5 adaptors, such as Rod1, Sna3 and Hua1 have been proposed to function later in the endocytic pathway, downstream of internalisation(Hovsepian et al., 2018; MacDonald et al., 2020, 2012b) these adaptors may be more important for downregulation when glucose is replaced with an alternative carbon source.

Additionally, the Cos proteins, which are also upregulated during nutritional stress and function late in the endosomal process, have been shown to help usher cargoes to MVBs for degradation (MacDonald et al., 2015a; MacDonald et al., 2015b). A similar mechanism controlling both degradation and recycling pathways has been described for nitrogen related starvation and TORC1 signalling. Cellular amino acid sufficiency is relayed to TORC1 via the Rag GTPases (Powis and Virgilio, 2016) and TORC1 inhibition following amino acid withdrawal results in an increase in Rsp5-adaptor complex activity, in a phosphoregulated pathway involving Npr1 (MacGurn et al., 2011). This increase in ubiquitin-mediated degradation in response to amino acid starvation coincides with a decrease in recycling back to the surface. Recycling in inhibited during Leucine starvation, which is sensed by the Rag GTPases that are required for efficient recycling back to the surface, in a process that relies on the co-factor Ltv1 (MacDonald and Piper, 2017).

One initially contrary finding is that eisosomes, which antagonise endocytosis (Gournas et al., 2018; Grossmann et al., 2008; Moharir et al., 2018; Spira et al., 2012), contain many factors that are transcriptionally upregulated in raffinose (**Figure 6B**), suggesting surface cargoes are controlled within subdomains of the plasma membrane itself. This finding can be rationalised by recent models that propose cargo protection in response to nutritional stress retains surface proteins for later use (Gournas et al., 2018; Moharir et al., 2018). Similarly, although our data shows levels of surface proteins are largely endocytosed and degraded in response to raffinose, there is a small population, for example Mup1-GFP (**Figure 6C**), that are retained and concentrated within eisosomes in these conditions. Increased Pil1 intensity during in raffinose, in addition to our 4D confocal microscopy capturing structural rearrangements of Pil1-mCherry at eisosomes that most significantly trap Mup1-GFP (**Supplemental Video 2**), suggests significant conformational changes occur in response to raffinose, increasing in size and possibly deepening to better sequester cargo. This also explains the initially confounding result that the diffusion coefficients from our single molecule analysis of Pil1-mCherry increased upon raffinose treatment (**Figure S6e**). Although single-cell stoichiometry estimates for Pil1 showed an increase in raffinose, the single-eisosome analysis revealed most of these changes occurred from eisosomes that retain cargo. Although nitrogen starvation and the broad nutritional stress of incubating cells for 12 hours after they reach stationary phase results in an increase in eisosomal numbers (Gournas et al., 2018), we did not observe an increase in eisosomal number in raffinose. This may be explained by the relatively short incubations in raffinose (30 – 120 minutes) necessary to observe the cargo downregulation and eisosome sequestration phenotypes documented in this study. As removal of carbon source results in a decrease in Nce102 at eisosomes (Appadurai et al., 2019), we predict that raffinose treatment, which increases transcript levels of *NCE102* around 6-fold, is triggering a distinct response to specifically upregulate eisosomes to retain nutrient transporters during the metabolic challenge of suboptimal sugars. A central finding of our study is that the sequestration of amino acid transporters provides a physiological benefit to cells during changes in sugar availability, by protecting a reservoir of nutrient transporters during glucose limited conditions. Upon a return to more favourable nutritional conditions, wild-type cells grow better than mutants that fail to properly sequester cargo in eisosomes (**Figure 7G**).

The finding that *YAP1801*, *YAP1802*, and *GPA2* are regulated in response to glucose via Mig1/Mig2 shows that membrane trafficking factors, in addition to genes centred around sugar metabolism, are modulated to adapt to changes in sugar availability. We did note that the levels of *Yap1801/02* and *GPA2* were elevated more when cells were acutely shifted to raffinose than when wild-type cells were compared with *mig1*Δ *mig2*Δ mutants. This might be explained by additional transcriptional regulators contributing to the glucose starvation response or that *mig1*Δ *mig2*Δ mutants have acquired additional mutations to compensate for the systemic loss of Mig1/Mig2. As gene deletions are often associated with secondary mutations, many of which affect stress responses (Teng et al., 2013), this latter idea is reasonable for *mig1*Δ cells, which are defective in various processes (Santangelo, 2006). The reduced effect in *mig1*Δ *mig2*Δ cells when compared to raffinose might also explain why we do not observe any changes in *PIL1* transcript levels in *mig1*Δ *mig2*Δ cells, as there was only a modest increase in raffinose (~2 fold). That said, as the other eisosomal genes are sensitive to raffinose treatment and do not contain Mig1 consensus binding sites, further supports the notion other transcriptional regulators must also be involved. This transcriptional response, like the modulation of internalisation and recycling described above, allows for acute metabolic control during in response to fluctuating nutrient conditions. Many elements of this control involve proteins highly conserved throughout evolution, so we propose similar regulation of these pathways and membrane subdomains in other eukaryotes could maintain metabolism in response to changes in extracellular nutrients.

## METHODS

### Reagents

Plasmids (Supplemental Table S2) and yeast strains (Supplemental Table S3) used in this study are listed in supplementary information. Polyclonal antibodies raised against GFP (Urbanowski and Piper, 1999), Rsp5 (Stamenova et al., 2004) and monoclonal Glyceraldehyde 3-phosphate dehydrogenase / GAPDH (clone 6C5; Ambion) and carboxypeptidase Y / CPY (clone 10A5-B5; Molecular Probes) antibodies, were used for immunoblot analysis.

### Cell culture

Yeast were grown in rich yeast extract peptone dextrose (YPD) media (2% glucose, 2% peptone, 1% yeast extract) or synthetic complete (SC) minimal medium (2% glucose, yeast nitrogen base supplemented with appropriate amino acid and base dropout mixtures; Formedium, Norfolk, UK) for maintaining of plasmids. Cells were routinely grown overnight to early / mid-log phase log phase (OD_600_ =<1.0) prior to experimental procedures, unless otherwise stated. For glucose starvtion experiments, rich and minimal media were supplemented with 2% raffinose instead of glucose but were otherwise identical. Geneticin (Formedium), used at a concentration of 250 μg/ml in rich media, and methotrexate (Alfa Aesar), used at a working concentration of 20 mM, was prepared in SC minimal media as described (MacDonald and Piper, 2015). Expression of proteins from the *CUP1* promoter was achieved by addition of 50 - 100 μM CuCl2 to the media for at least 1 hour prior to downstream analysis.

### Confocal microscopy

Cell were grown to mid-log phase and prepared for fluorescence microscopy experiments by concentration in minimal media before imaging or from a slide or glass-bottom dish using Laser scanning confocal microscopes (LSM710 or LSM880 equipped with an Airyscan module; Zeiss) with a 63x Differential Interference Contrast (DIC) objective with a 1.4 numerical aperture. Argon laser excitation, 488 nm with emission filter set to 495 – 550 nm (for GFP, mGFP and mNeonGreen) and 561 nm with 570 – 620 nm emission filter (for mCherry and FM4-64) were used.

### Microfluidics & time lapse microscopy

Cells were grown to early log phase (approximately OD_600_ = 0.2) overnight and then adhered to 35 mm concavalin A coated glass bottom dishes (Ibidi GmbH, Germany) prior to live cell imaging at room temperature. Plates were prepared by adding 1 mg/mL concavalin A in water to coverslip for 5 minutes prior to several wash steps; plates were routinely stored at 4°C. Sterile media exchanges were performed using syringes through tubing fused to the lid of 35 mm dishes. Live cells were labelled with 0.8 μM FM4-64 using microfluidics for 2 - 5 minutes before flushing with media containing 2.4 μM SCAS (Biotium, Hayward, CA) for 30 seconds at room temperature to quench extracellular dye before further imaging.

### Image analysis

Micrographs were processed by Zeiss Zen and Fiji software. For time lapse videos bleach correction carried out using histogram matching method (https://imagej.net/Bleach_Correction). For images of the top section of a yeast cell 5 slices with 0.18m spacing are combined using the average intensity of the maximum intensity z-projection. Movies made using time lapse image, of varying frame time points, all indicated in the movies. Numbers of eisosomes were identified using Tracking software written in MATLAB and reported previously (Miller et al., 2015; Wollman et al., 2017; Wollman and Leake, 2015).

### Slimfield microscopy and fluorescent foci analysis

Slimfield excitation was implemented via co-aligned 488 nm and 561 nm wavelength 50 mW lasers (OBIS, Coherent) de-expanded to direct a beam onto the sample full width at half maximum of ~30μm. For visualisation of both fluorophores we employed rapid Alternating Laser Excitation (ALEX) with 5 ms exposure time per image frame for each laser (Syeda et al., 2019). Fluorescence emission was captured by a 1.49 NA oil immersion objective lens (Nikon), and subsequently split into separate green and red detection channels using a dual-pass green/red dichroic mirror centred at long-pass wavelength 560nm combined with 25 nm bandwidth emission filters (Chroma) centred on 525 nm and 594 nm wavelengths. Each channel was imaged separately at 5 ms exposure time by an EMCCD camera Prime 95B^TM^ Scientific CMOS, Teledyne Photometrics) using 50 nm/pixel magnification. The focal plane was set to mid-cell height using the bright field appearance of cells. As photobleaching of mGFP and mCherry proceeded during Slimfield excitation distinct fluorescent foci could be observed of width 250-300 nm, consistent with the diffraction-limited point spread function of our microscope system, which were tracked and characterized in terms of their stoichiometry and apparent microscopic diffusion coefficient. Distinct fluorescent foci that were detected within the microscope’s depth of field could be tracked for up to several hundred ms, to a super-resolution lateral precision σ = 40 nm using a bespoke single particle tracking algorithm. The molecular stoichiometry for each track was determined by dividing the summed pixel intensity values associated with the initial unbleached brightness of each foci by the brightness corresponding to that calculated for a single fluorescent protein molecule (either mGFP for 488 nm wavelength excitation or mCherry for 561 nm wavelength excitation) measured using a step-wise photobleaching technique described previously (Shashkova et al., 2018; Wollman et al., 2019). The apparent microscopic diffusion coefficient D was determined for each track by calculating the initial gradient of the relation between the mean square displacement with respect to tracking time interval using the first five time intervals values while constraining the linear fit to pass through 4σ^2^ on the vertical axis corresponding to a time interval value of zero. Cell body and membrane boundaries were detected based on the mCherry fluorescence images, considering the membrane width as 7nm.

### Gene expression (qPCR) analysis

Yeast cultures (10 ml) were grown overnight to midlog phase in YPD or exchanged with YPR for indicated times prior to spheroplasting in yeast lysis buffer (1 M sorbitol 100 mM EDTA, 0.1% β-mercaptoethanol) with 5μl Zymolase added for 2 minutes at room temperature. Extraction of total RNA media was performed using an RNeasy kit (QIAGEN) protocol with an additional DNaseI treatment, followed by a second DNaseI digestion with the TURBO DNA-free kit (Invirogen). Extracted mRNA was then used for cDNA synthesis with the SuperScript IV reverse transcriptase (Invitrogen). 50ng/μ random hexamers, and 10mM dNTPs were added to 5μg RNA. Samples were incubated at 65°C for 5 minutes before addition of 100mM DTT, Ribonuclease inhibitor and the Superscript IV reverse transcriptase to initiate the reaction (10 mins 23°C; 10 mins 55°C; 10 mins 80°C) and then used immediately for the qPCR reaction. For qPCR optimisation, two sets of primers targeting amplicons 70 – 170 bp in length were designed for each gene of interest, and the best performing was chosen for use in quantitation (Supplemental Table S2 for oligonucleotide sequences). Single product amplification was confirmed through PCR using genomic DNA, and near-100% amplification efficiencies were confirmed through qPCR on a standard curve of known input quantity, all performed in duplicate reactions (examples shown in Supplemental Figure s2). All qPCR reactions were performed on 20 ng cDNA, or relevant negative controls, in 20 μl reactions containing 350 nM forward and reverse primers, and 10 μl Fast SYBR™ Green Master Mix (ThermoFisher Scientific). Reactions were carried out using the QuantStudio 3 system (Thermofisher) under the following conditions: 40 cycles of 95°C for 1 second, 20 seconds 60°C, before a continuous ramp from 60°C to 95°C at a rate of 0.1 °C/S for melt curve analysis. Gene expression levels were quantified using the comparative Ct (ΔΔCt) method, relative to the expression of the housekeeping gene (*ACT1* or *HEM2*) and normalized to control sample (e.g. wild type when testing mutant strains; glucose when assessing effects of raffinose time-course). To avoid bias caused by averaging data that has been transformed through the equation 2^-Ct^ to give foldchanges in gene expression, all statistics were performed on Ct, values.

### Immunoblotting

Equivalent amounts of cells were harvested from early and late log phase (OD_600_ = 0.5 and 2.0) yeast cultures. Lysates were generated by alkali treatment with 0.2 M NaOH for 3 minutes prior resuspension in lysis buffer (8 M Urea, 10% glycerol, 50mM Tris.HCl pH 6.8, 5% SDS, 0.1% bromophenol blue, 10% 2-Mercaptoethanol). Proteins were resolved by SDS-PAGE prior to transfer to a nitrocellulose membrane using the iBlot 2 dry transfer system (ThermoFisher Scientific). Membrane was probed with appropriate antibodies and visualised by Enhanced chemiluminescence (ECL) super signal Pico Plus (ThermoFisher Scientific), with intensity digitally captured using a ChemiDoc Imager (BioRad).

### FM4-64 efflux assay

FM4-64 efflux was measured in cells grown in SC 2%glucose media to late-log phase overnight. 1ml of each sample was pelleted and resuspended in YPD for 3 hours. 1ml of cells (OD = 1.0) was harvested and resuspended in 100 μl YPD containing 40 μM FM4-64. Cells were incubated for 8 minutes at room temperature then harvested by washing 3 times in ice cold SC 2% glucose media for 3 minutes per wash. Cells were resuspended in 100 μl SC 2% glucose media and stored on ice before addition of room temperature glucose or raffinose media and fluorescence intensity measured by flow cytometry using a LSRFortessa instrument (BD Biosciences) with excitation 561nm, laser emission filter 710 / 50.

### Flow cytometry

Intensity from GFP expressing (excitation laser 488 nm, emission filter (525 / 40 nm) and FM4-64 or mCherry (561 nm excitation laser, 710 / 50 nm emission filter) labelled live cells at room temperature was recorded using a CytoFLEX flow cytometer (Beckman Coulter) and intensity measurements from gated cells measured with FCS Express v6.0 (DeNovo software). Typically, 100,000 cells, gated for fluorescence positive yeast cells (forward/side scatter), were flowed at ~600 V to maintain a rate of approximately 500 – 1000 cells measured per second. Unlabelled / non-expressing cells were used for background calibration. Channel recordings from other detectors (e.g. 530 / 50 nm during FM4-64 acquisitions) were also recorded to measure background autofluorescence. Intensity measurements are plotted. The co-efficient of variation (CV) equal to the population standard deviation divided by the population mean is expressed as a percentage.

### Yeast growth and recovery assays

Yeast cultures were grown overnight in glucose containing YPD or SC media diluted to OD_600_ = 0.4 and used to create a 10-fold serial dilution in a sterile 96-well plate, which were then plated on indicated media, grown for 2 days at 30°C before growth was recorded using a Phenobooth Imager (Singer Instruments). For recovery-based assays, cells were exchanged to raffinose media for 2 hours prior to equivalent cell numbers were harvested and either plated on indicated solid media or brought up in glucose containing minimal media containing various concentrations of uracil and OD_600_ measurements recorded every 60 minutes using a MultiSkanGo plate reader and SkanIt software (ThermoFisher Scientific).

### Statistical analyses

The statistical significance for each experimental condition (e.g. raffinose media, mutant yeast strains) in comparison to control conditions (e.g. glucose media, wild-type cells) was calculated using unpaired t-test / Holm-Sidak method in GraphPad Prism v8.3.1. An asterisk (*) is used to denote p-values of <0.05 in graphs but all generated p-values for each experiment are included in Supplemental Table S5.

## Supporting information

Supplemental Figures, legends and references

## ACKNOWLEDGMENTS

We are grateful to the York Bioscience Technology Facility, including Peter O’Toole and the imaging team for technical assistance, Karen Hogg for help with flow cytometry, and Sally James for guidance with gene expression analysis. We would like to thank Hans Ronne for the generous gift of *mig1*Δ *mig2*Δ and *mig1*Δ *mig2*Δ *mig3*Δ yeast strains, which were used for initial work and to David Teis for fluorescently labelled Vps4 strains. We are grateful to Chris Burd for suggestions based on an earlier version of this story and Nia Bryant for helpful comments on the manuscript. This research was supported by a Sir Henry Dale Research Fellowship from the Wellcome Trust and the Royal Society 204636/Z/16/Z (CM), and EPSRC EP/T002166/1 (ML), BBSRC BB/R001235/1 (ML), Marie Curie EU FP7 ITN Ref 764591 (ML), and the Leverhulme Trust RPG-2019-156/ RPG-2017-340 (ML).

## DECLARATION OF INTERESTS

The authors declare no competing interests.

