## Supplemental Figures, legends and references for "A glucose starvation response governs endocytic trafficking and eisosomal retention of surface cargoes"

### SUPPLEMENTAL FIGURES AND LEGENDS

- Figure S1 – Supplement to Figure 1
  ***Differential cargo trafficking effects following glucose starvation***

- Figure S2 – Supplement to Figure 2
  **qPCR optimisation and time-lapse microscopy of Mig2-GFP**

- Figure S3 – Supplement to Figure 3
  **Localisation of Yap1801 and Yap1802**

- Figure S5 – Supplement to Figure 5
  **Ste3-GFP-DUb and Galpha subunits localisations**

- Figure S6 – Supplement to Figure 6
  **Analyses of eisosomes in response to changes in glucose levels**

- Figure S7 – Supplement to Figure 7
  **Growth and cargo localisation defects in eisosome mutants**

### SUPPLEMENTAL VIDEOS

- Supplemental Video S1
  **Mig1-GFP translocation is sensitive to extracellular carbon source**

- Supplemental Table S2
  **Mup1-GFP and Pil1-mCherry colocalise following glucose starvation**

### SUPPLEMENTAL TABLES

- Supplemental Table S1
  **Functional clustering of Mig1 candidates**

- Supplemental Table S2
  **qPCR primers**

- Supplemental Table S3
  **Yeast Strains used in this study**

- Supplemental Table S4
  **Plasmids used in this study**

- Supplemental Table S5
  **Statistical analyses**

### SUPPLEMENTAL REFERENCES

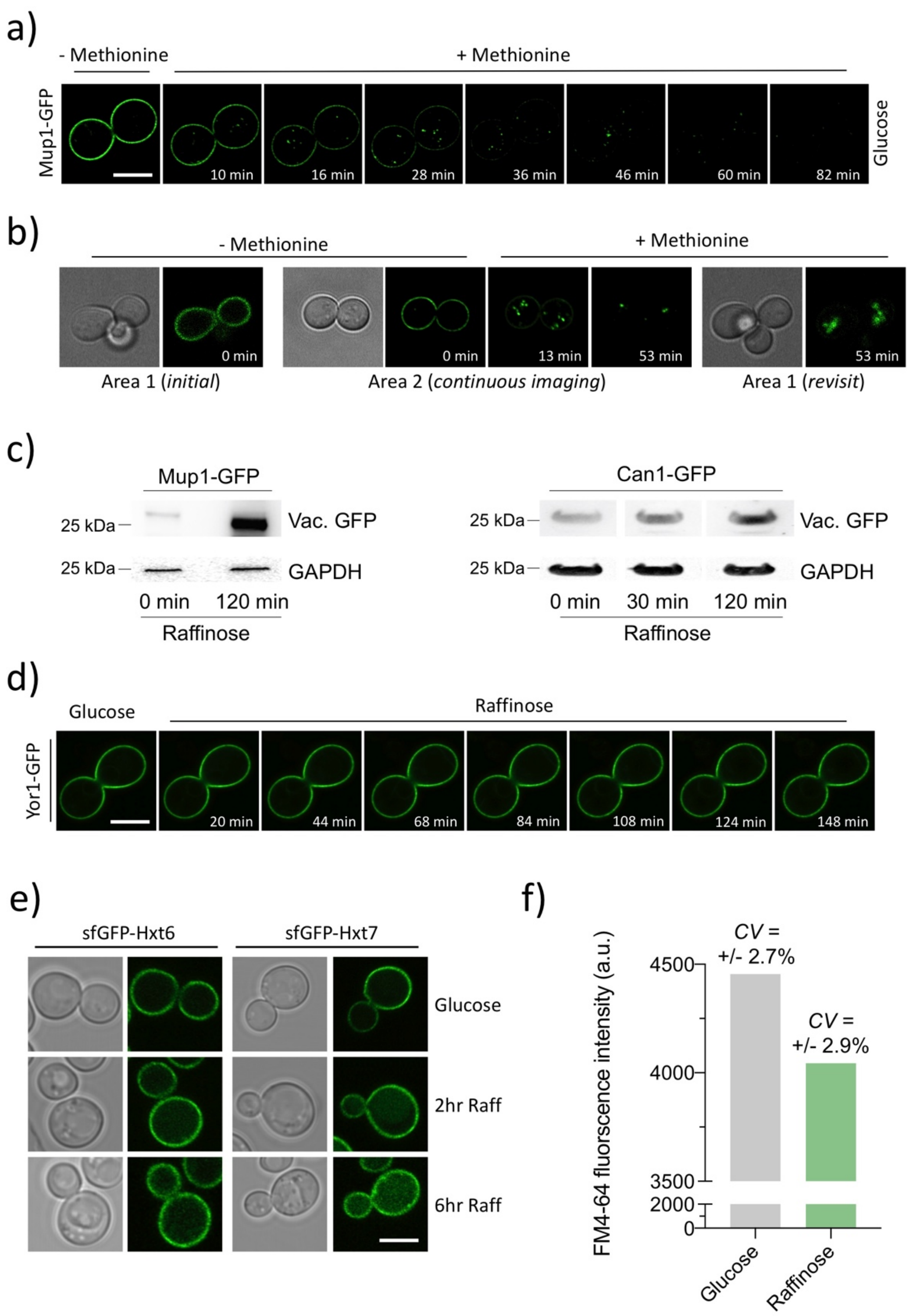


*Figure S1: Differential cargo trafficking effects following glucose starvation.* ***a)*** *Wild-type cells expressing Mup1-GFP were grown to mid-log phase in SC media lacking methionine, processed for time-lapse microscopy and then imaged every 2 minutes following addition of 20 µg/ml methionine.* ***b)*** *Vacuolar GFP bleaching in wild-type cells expressing Mup1-GFP grown to mid-log phase in SC media lacking methionine and processed for time-lapse microscopy. Area 1 was imaged before addition of methionine, before moving to a distinct region of the same plate (Area 2) for continuous imagining from 0 – 53 minutes of methionine addition. Following this period, Area1 was re-visited and imaged to show the difference in photobleaching of vacuolar sorted Mup1-GFP.* ***c)*** *Levels of vacuolar processed GFP in cargo Mup1-GFP (left) and Can1-GFP (right) expressing cells were assessed in glucose and raffinose treated cells by immunoblotting lysates with GFP antibodies. Loading was assessed with anti-GAPDH antibodies.* ***d)*** *Wild-type cells expressing Yor1-GFP from the CUP1 promoter buy addition of 50µM copper chloride were grown to mid-log phase in glucose containing media, processed for time-lapse microscope and imaged for indicated time course after exchange with raffinose media.* ***e)*** *Strains expressing GFP tagged Hxt6 and Hxt7 expressed from the NOP1 promoter were grown in glucose or raffinose media for indicated periods prior to fluorescence microscopy. f) Wild-type cells grown either in glucose media or exchanged with raffinose for 15 minutes were incubated with YPD containing 40 µM FM4-64 dye for 4 minutes at room temperature before ice cold washes were performed with minimal media to remove excess dye. Mean fluorescence of ~10,000 cells was then measured by flow cytometry. Scale bar, 5 µM.*

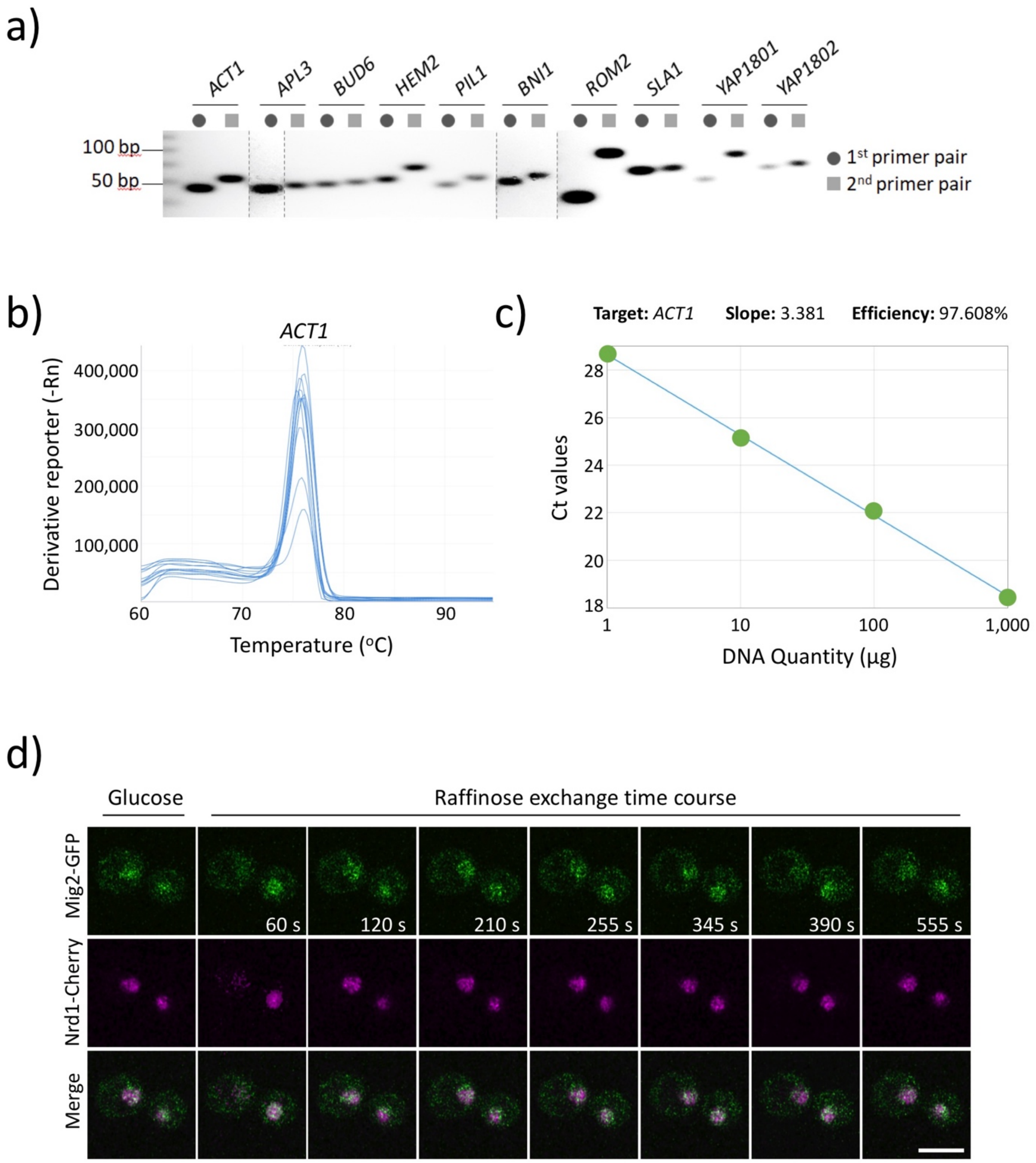


*Figure S2: qPCR optimisation and time-lapse microscopy of Mig2-GFP:* ***a)*** *Indicated oligo pairs were used to generate PCR products from a gDNA template and analysed by agarose gel of primer pair validation of the qPCR using wild-type cells mRNA.* ***b)*** *Melt curve analyses for all qPCR primer pairs were carried out to ensure no primer dimer species were detected, read out shows example for ACT1.* ***c)*** *Primer pair efficiency was also performed for all primer pairs used in this study (shown for ACT1), with ΔCt values across a 10-fold serial dilution plotted (slope = -3.33 equivalent to 100% efficiency). Data for all primer pair analysis is recorded in Supplemental Table S2.* ***d)*** *Wild-type cells expressing Mig2-GFP and Nrd1-mCherry were grown to mid log phase in minimal media prior to processed for time-lapse microscopy. Images were captured every 5 seconds following raffinose exchange, with representative time-slices shown. Scale bar, 5 μM.*

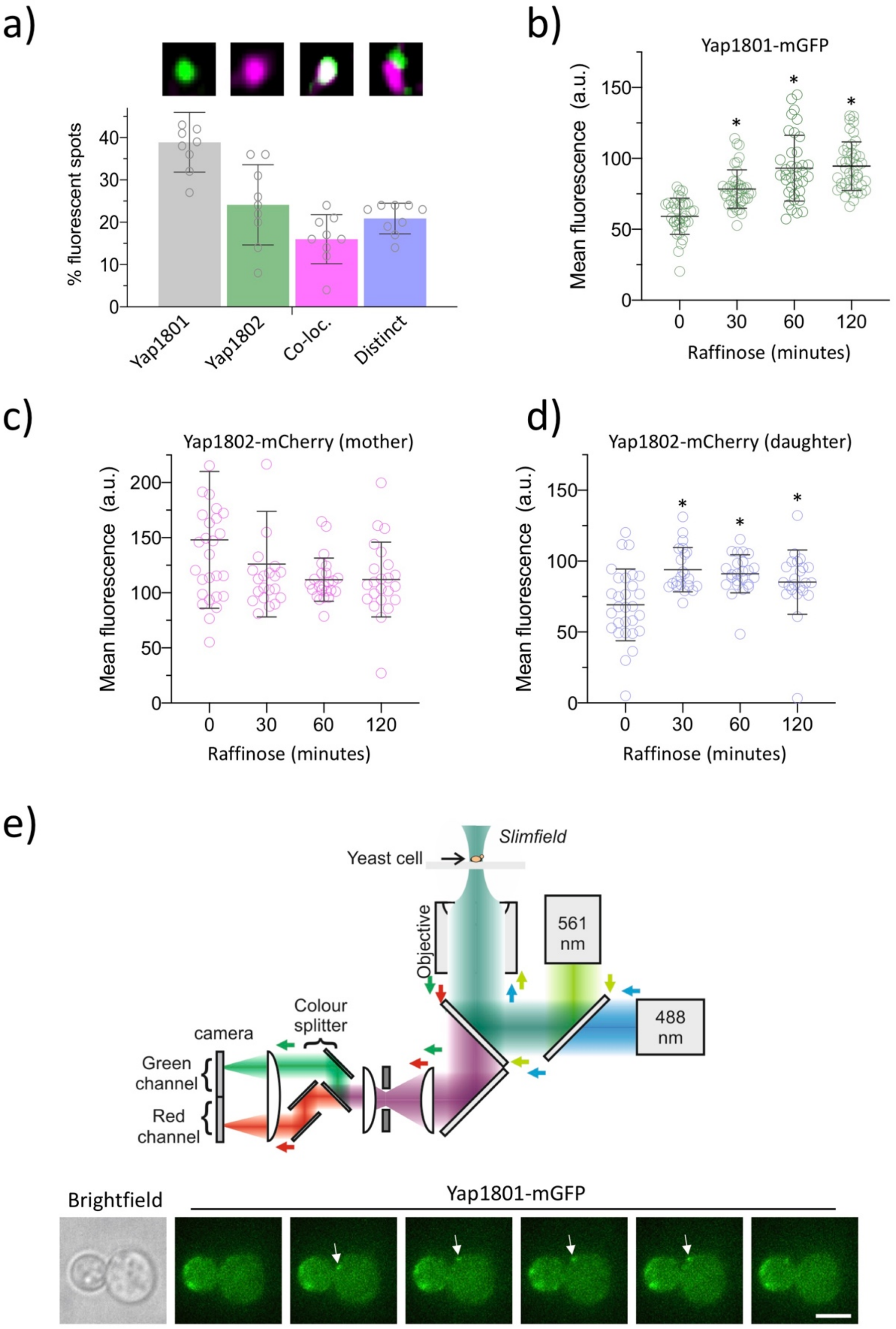


*Figure S3: Localisation of Yap1801 and Yap1802:* ***a)*** *Histogram showing the co-localisation analysis of Airyscan confocal images of Yap1801-mGFP and Yap1802-mCherry expressing wild-type cells grown to mid-log phase in SC selective media. Error bars showing standard deviation (n = 235 foci analysed). Examples of each localisation category is shown (upper).* ***b)*** *Histogram showing average Yap1801-mCherry (total cell) and Yap1802-mCherry (just mother cell) intensity in glucose and raffinose media. Intensity was averaged from n=>36 cells per condition over 3 biological replicates, with error bars showing standard deviation.* ***c)*** *Histogram showing the mean fluorescence of wild-type cells expressing Yap1802-mCherry (daughter cell) when grown in glucose or raffinose containing media for indicated time course over 3 biological replicates, with error bars showing standard deviation.* ***d)*** *Slimfield microscopy, schematic diagram showing set-up for dual-colour imaging of yeast cells. Lower panels show Yap1801-mGFP fluorescent spot (white arrow) tracking from images acquired every 5ms. * indicates t-test p-values <0.05. Scale bar, 5 μM.*

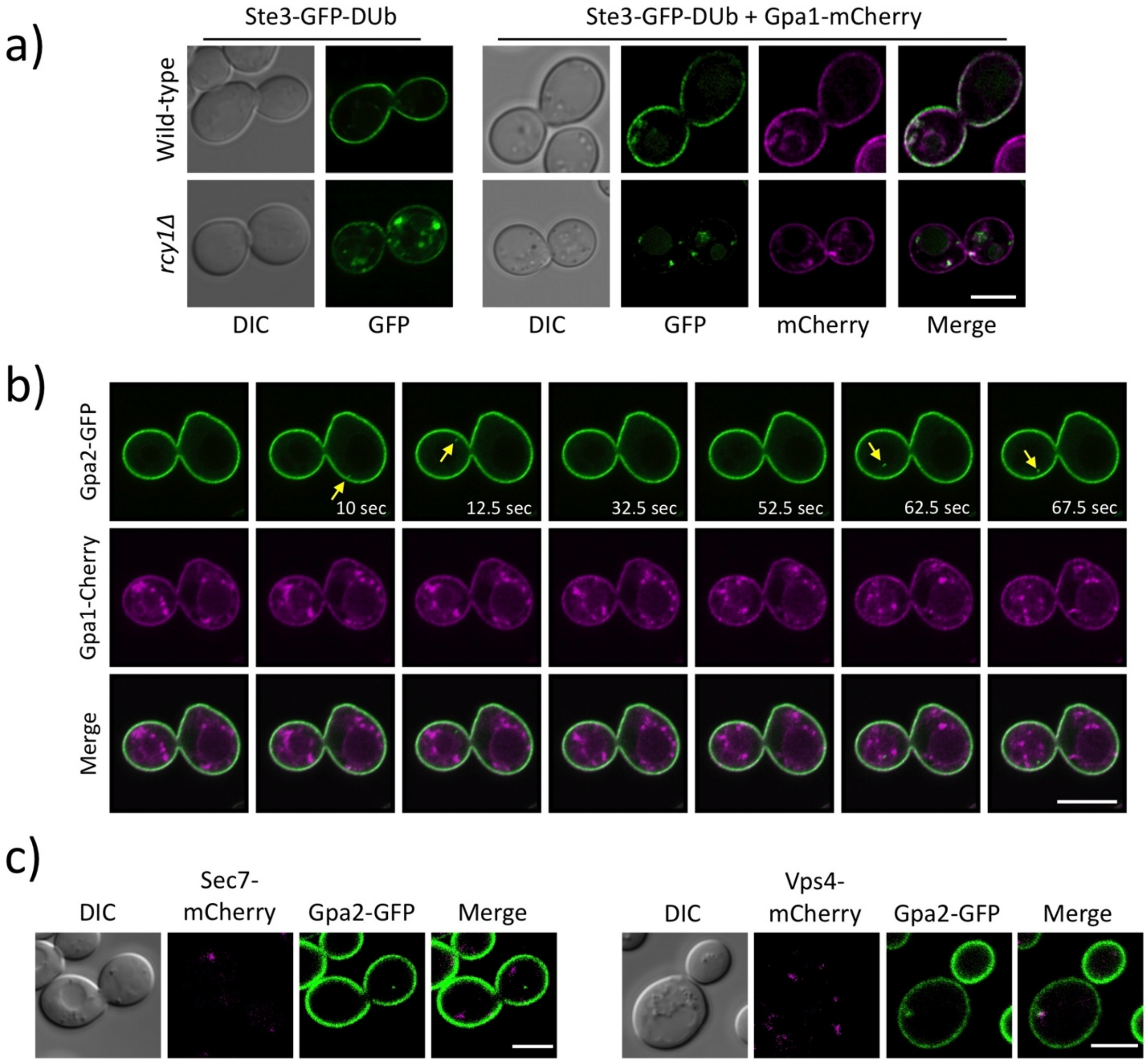


*Figure S5: Ste3-GFP-DUb and Galpha subunits localisations:* ***a)*** *Wild-type and rcy1Δ cells stably expressing Ste3-GFP-DUb were imaged alone (left) and when co-expressing Gpa1-mCherry, induced under the control of the CUP1 promoter using 50 μM copper chloride in the media (right).* ***b)*** *Time series of wild-type cells expressing Gpa2-GFP and Gpa1-GFP from the CUP1 promoter in the presence of 50 μM copper chloride. Cells were grown to mid-log phase and prepared for confocal imaging. Cells are imaged every 2.5 seconds; yellow arrow denotes intracellular foci of Gpa2-GFP.* ***c)*** *Wild-type cells stably expressing either Sec7-mCherry (left) or Vps4-mCherry (right) were transformed with a plasmid expressing Gpa2-GFP under the CUP1 promoter in the presence of 50 μM copper chloride grown to mid-log phase in SC media before preparation for confocal analysis.*

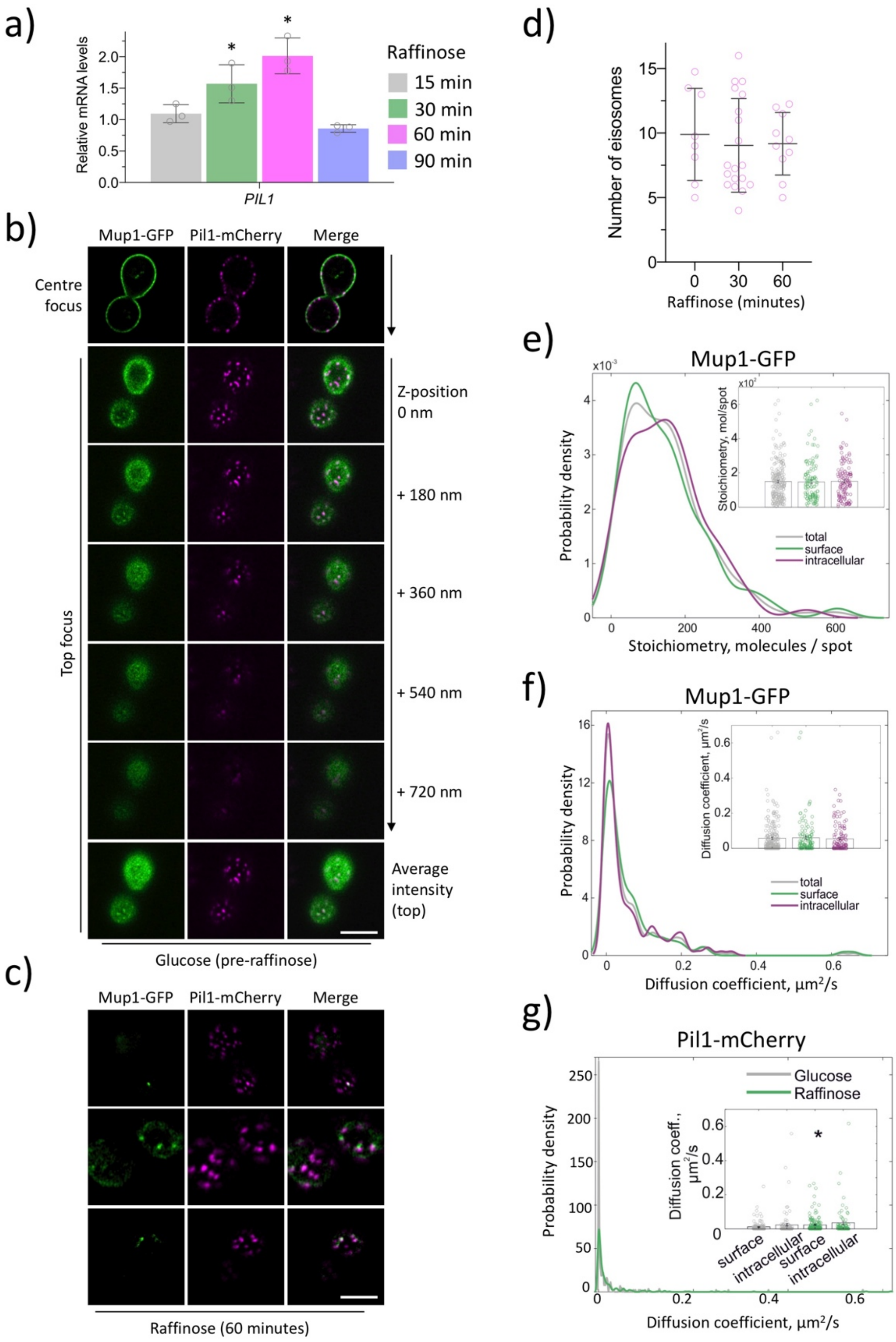


*Figure S6: Analyses of eisosomes in response to changes in glucose levels:* ***a)*** *Quantitative RT-PCR of PIL1 performed from RNA extracted from wild-type cells grown in glucose media and relative levels compared to cells grown in raffinose media for indicated time course. Error bars show the standard deviation from 3 biological replicates (each averaged from 3 technical replicates).* ***b)*** *Wild-type cells expressing Pil1-mCherry and Mup1-GFP were grown to mid-log phase in SC media and prepared for confocal imaging. Cells were focussed to the top of the cell and imaged at 0.18[illegible]m intervals in the z-axis. An average intensity projection from the indicated 5 slices is shown (lower panel).* ***c)*** *Number of Pil1-mCherry marked eisosomes per cell in glucose (n = 33 cells) and 30 minutes raffinose (n = 71 cells) and depicted with standard deviation error bars.* ***d)*** *Kernel density plots of Mup1-GFP stoichiometry (top) and diffusion coefficients (bottom) distribution of fluorescent foci in the whole cell (grey), on the surface (green) or inside the cell (purple). Inserts: jitter plots of stoichiometries and diffusion coefficients of fluorescent foci detected in the whole cell, on the surface or inside the cell. Error bars represent standard error.* ***e)*** *Kernel density plot of Pil1-mCherry foci diffusion coefficients distribution in cells incubated in glucose (grey) or raffinose (green). Insert: jitter plot of diffusion coefficients of fluorescent spots identified on the surface and inside the cell in yeast cultures incubated in glucose (grey) and raffinose (green). Error bars represent standard error. * indicates t-test p-values <0.05. Scale bar, 5 μm.*

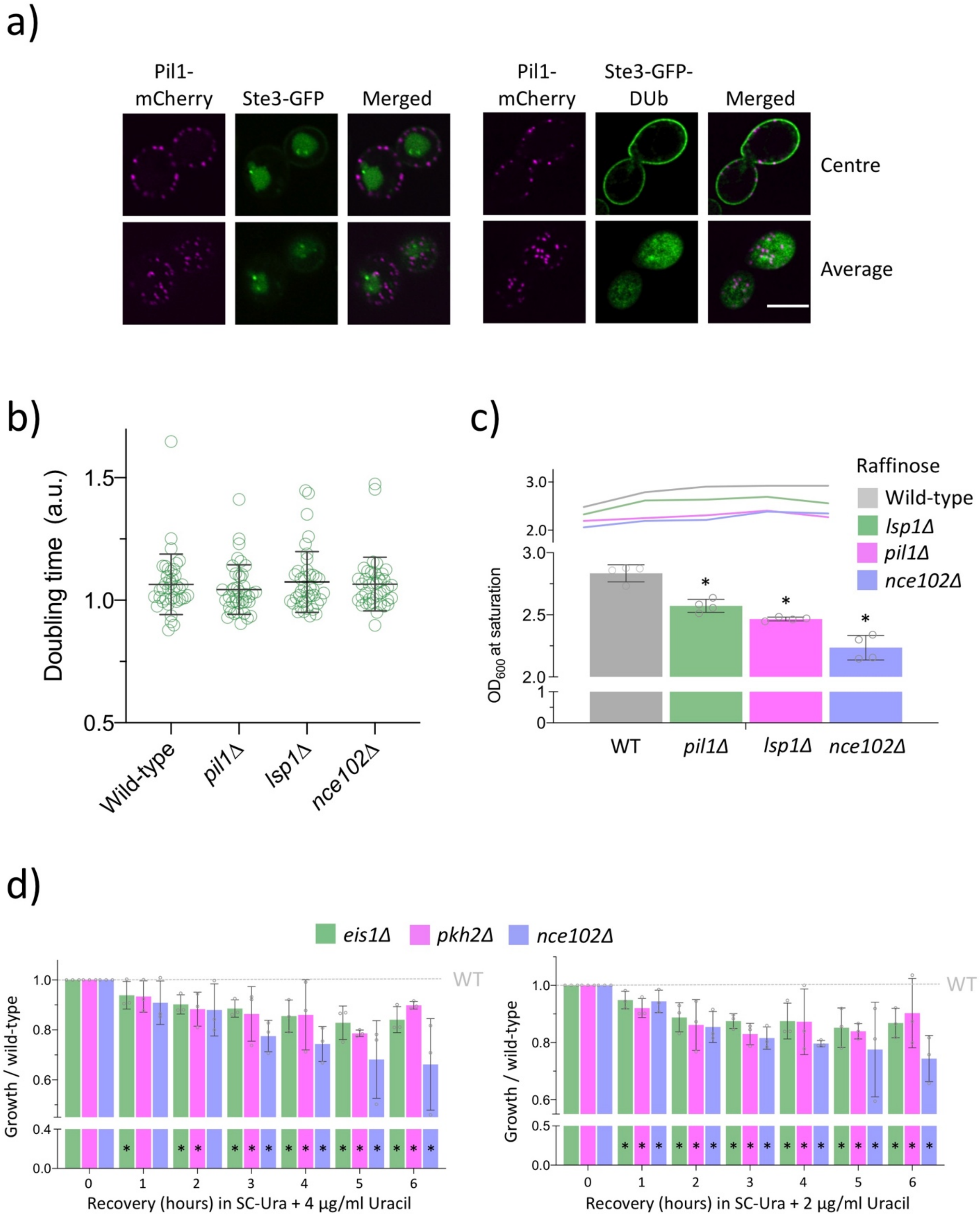


*Figure S7: Growth and cargo localisation defects in eisosome mutants.* ***a)*** *Wild-type cells co-expressing Pil1-mCherry and either Ste3-GFP (left) or Ste3-GFP-DUb (right) were imaged by 3D confocal Airyscan.* ***b)*** *Triplicate cultures were grown to mid-log phase overnight, diluted to low optical density and then grown in 5ml cultures. Samples were taken for OD$_{600}$ measurements every hour over 8 – 10 hours and used to calculate average doubling times, with distribution displayed in scatter plot.* ***c)*** *Indicated strains were grown to mid-log phase overnight and allowed to reach saturation, with OD$_{600}$ measurements captured every hour, depicted as a line graph (upper), or the maximum OD$_{600}$ shown as a histogram (lower).* ***d)*** *Indicated mutants were grown to mid-log phase before incubation in raffinose media for two hours. Cells were then resuspended in SC media containing glucose and 0.5µg/ml Uracil and OD$_{600}$ measurements recorded over 6-hour recovery period, depicted as histogram. Error bars show standard deviation from 3 biological replicates. * indicates t-test p-values <0.05. Scale bare, 5 µm.*
